# A high-resolution map of functional miR-181 response elements in the thymus reveals the role of coding sequence targeting and an alternative seed match

**DOI:** 10.1101/2023.09.08.556730

**Authors:** Nikita A. Verheyden, Melina Klostermann, Mirko Brüggemann, Hanna M. Steede, Anica Scholz, Shady Amr, Chiara Lichtenthaeler, Christian Münch, Tobias Schmid, Kathi Zarnack, Andreas Krueger

## Abstract

MicroRNAs (miRNAs) are critical post-transcriptional regulators in many biological processes. They act by guiding RNA-induced silencing complexes to miRNA response elements (MREs) in target mRNAs, inducing translational inhibition and/or mRNA degradation. Functional MREs are expected to predominantly occur in the 3’ untranslated region and involve perfect base-pairing of the miRNA seed. Here, we generate a high-resolution map of miR-181a/b-1 (miR-181) MREs to define the targeting rules of miR-181 in developing murine T-cells. By combining a multi-omics approach with computational high-resolution analyses, we uncover novel miR-181 targets and demonstrate that miR-181 acts predominantly through RNA destabilization. Importantly, we discover an alternative seed match and identify a distinct set of targets with repeat elements in the coding sequence which are targeted by miR-181 and mediate translational inhibition. In conclusion, deep profiling of MREs in primary cells is critical to expand physiologically relevant targetomes and establish context-dependent miRNA targeting rules.

**Key Points:** • Deep profiling identifies novel targets of miR-181 associated with global gene regulation.

• miR-181 MREs in repeat elements in the coding sequence act through translational inhibition.

• High-resolution analysis reveals an alternative seed match in functional MREs.

## Introduction

MicroRNAs (miRNAs) are small, noncoding RNAs that constitute a critical layer in the post-transcriptional regulation of gene expression. They abundantly occur in eukaryotic cells where they impact many biological processes, including immune system development and function (1). miRNAs typically span 20-24 nucleotides (nt) and are generated via multiple cleavage steps from miRNA precursors. They act by guiding Argonaute-containing RNA-induced silencing complexes (RISCs) to miRNA recognition elements (MREs) in their target mRNAs. Efficient targeting depends on a combination of biochemically established targeting rules and contextual factors of a given miRNA and cell type. In mammals, the main Argonaute protein is Ago2 which is the only Argonaute family member capable of directly cleaving RNA (2). Commonly, miRNA-loaded RISCs recruit exonucleases to rapidly degrade the targeted mRNA. Alternatively, RISC-associated factors may compete with translation initiation factors for cap binding or cause ribosome stalling, both resulting in translational inhibition (3–5).

Typically, a given miRNA can target multiple distinct mRNAs in any given cell, referred to as the miRNA targetome. Since the abundance of potential miRNA recognition elements (MREs) generally surpasses the concentration of the corresponding miRNA (6), hierarchical targeting of MREs is critical for any miRNA to elicit biological functions. Canonical targeting of a miRNA to an MRE primarily occurs through base-pairing between the 6 nt long miRNA seed region (positions 2-7) and complementary sequences in the 3’ untranslated region (3’UTR) of the target mRNA, referred to as seed match (7,8). Extended base-pairing at adjacent positions contributes to prioritization of miRNA binding to distinct MREs, with 8-mer (positions 1-8), 7-mer-A1 (positions 1-7) and 7-mer-m8 (positions 2-8) seed matches typically displaying the highest activity (9–11). In addition to seed extension, non-canonical base-pairing at 3’ positions of the miRNA may contribute to MRE binding and compensate for imperfect seed matches (12–15). Apart from these canonical or near-canonical MREs, alternative types of MREs have been described (12,13,16–18). However, the efficacy of 3’ supplemental pairing or non-canonical types of MREs remains unclear. Moreover, the hierarchies of miRNA–MRE binding and, concomitantly, repressive capacity are not universal, but may vary between miRNAs (11). Taken together, despite a number of global rules for predicting miRNA target selectivity, which have been implemented in many *in silico* tools, the prioritization of MREs for mediating RISC-dependent repression is dependent on the miRNA sequence and concentration as well as the overall pool of MREs present in a given cell types. The impact of a miRNA in a specific cellular context therefore remains hard to infer without targeted experiments.

The miRNA miR-181 constitutes a family of six miRNAs organized in clusters of miR-181a/b-1, miR-181a/b-2 and miR-181c/d in mice and humans. The generally more stable 5p strands of all six members share the same seed sequence ACAUUC – pairing to the seed match GAAUGU in the mRNA – but differ in length and sequence at the 3’ end. MiR-181 family members have been suggested to play critical regulatory roles in a large number of biological processes, including but not limited to the cardiovascular and immune systems (19). In the immune system, miR-181 is most prominently linked to the function and development of T cells (20). In peripheral T cells, levels of miR-181a decline with increasing age, resulting in reduced antiviral T-cell function in older individuals and mice (21–23). In developing T cells, miR-181a-1 and miR-181b-1 are among the most prominently expressed miRNAs (24–26). Deletion of the *Mirc14* locus encoding miR-181a/b-1 results in a profound defect in the development of unconventional T cells, including invariant Natural Killer T (iNKT) cells, Mucosal-Associated Invariant T (MAIT) cells, and regulatory T (Treg) cells (27–30). Mechanistically, this phenotype is consistent with miR-181 binding the transcripts of several negative regulators of T-cell receptor signaling, encoding for the phosphatases Ptpn22, Shp-2, Dusp6 and Dusp10, among others, resulting in lower overall signaling thresholds (31–33,20,30,24,34). Besides these well-characterized targets several hundred mRNAs have been proposed to be targeted by miR-181a/b and some of those, encoding Bcl-2, CD69, Pten, Nrarp or S1PR1 have been implicated in miR-181a/b-dependent regulation of T-cell development and function (34,30,24,35,32).

Here, we characterize the miR-181a/b targetome in primary murine thymocytes by deep profiling of MREs together with a genetic deletion of miR-181a/b-1. By combining chimeric Ago2 eCLIP and oligonucleotide-based miRNA enrichment with transcriptome and translatome sequencing, we find that miR-181 primarily acts through mRNA destabilization rather than translational inhibition. Repression of mRNAs correlates with increasing numbers of MREs which are primarily, but not exclusively, characterized by canonical seed binding and predominantly located in 3’UTRs. Surprisingly, we uncover a substantial fraction of MREs harboring an alternative seed match with similar repressive capacity. In addition, we characterize a specialized subset of MREs in satellite repeat elements, which act through translational inhibition in the coding region and suggest an evolutionary link between miR-181 regulation and transcription factor expansion in mammalian genomes. Taken together, our study sheds light on novel targeting networks in T-cell development and highlights the need for deep profiling and high-resolution analyses to decipher the rules governing context-dependent miRNA function.

## Materials and Methods Mice

MiR-181a/b-1^-/-^ mice (B6.*Mirc14*^tm1.1Ankr^) were generated as described in (29,36) and bred at Goethe University Frankfurt and Justus Liebig University Giessen. Experiments were performed using 3 miR-181a/b-1^-/-^ mice and 3 WT controls (C57BL/6) for RNAseq and ribosome footprinting. 3 miR-181a/b-1^-/-^ mice and 3 WT controls (C57BL/6) were used for eCLIP analysis. All mice were male and 6 weeks of age. The study was approved under permits FU/1255, Regierungspräsidium Darmstadt (for Goethe University Frankfurt) and 818_M (for Justus Liebig University Gießen).

### eCLIP

Total Thymocytes were isolated from 3 WT and 3 miR-181a/b-1^-/-^ mice and processed individually to obtain biological replicates. Isolated cells were then crosslinked using a using 254nm UV-C Mercury bulb in the (Analytik Jena CL-1000M). Cells were sent to Eclipse Bioinnovations (San Diego, CA, USA), where their miR-eCLIP^TM^ was performed with enrichment for mmu-miR-181a-1 (miRbase ID: MI0000697) and mmu-miR-181b-1 (miRbase ID: MI0000723) (37).

### RNA sequencing (RNAseq) and ribosome footprinting (RF)

Ribosome profiling was performed according to Ingolia et al. following the protocol below (36).

#### Animals and sample collection

We obtained thymi from 3 wild-type (WT) and 3 miR-181a/b-1^-/-^ knockout (KO) mice processed individually to obtain biological replicates. The thymi were extracted and immediately submerged in RPMI media with 0.1 mg/ml cycloheximide.

#### Cell extraction and splitting

Total thymocytes were extracted using a 70 µl mesh, while constantly being submerged in DPBS containing 0.1 mg/ml cycloheximide. The resulting samples were split into RF and RNAseq samples.

#### RF sample processing

For the RPF samples, we used the TruSeq Ribo Profile kit (Illumina) to process them according to the manufacturer’s instructions. Samples were digested with the TruSeq Ribo Profile Nuclease for exactly 45 min at room temperature, then purified using the RNA Clean & Concentrator-25 kit. RNA was quantified by NanoDrop (Thermo Fisher Scientific).

To remove rRNA, we used the magnetic beads from the Illumina Ribo-Zero Plus rRNA Depletion Kit and 5 µg of RNA following the manufacturer’s instructions. The total RNA and RF samples were purified using a modified RNA Clean & Concentrator-5 kit method (Zymo Research). The volume of each sample was increased to 100 µl, and 100 µl RNA Binding Buffer was added to the total RNA samples and 200 µl to the RF samples, along with 100 µl of 100% ethanol to the total RNA samples and 450 µl to the RF samples. After following the manufacturer’s protocol, RNA samples were eluted in 21 µl and RF samples in 11 µl.

RF samples were cleaned up using a 15% urea-polyacrylamide gel following the manufacturer’s protocol.

#### Total RNA sample processing

For the total RNA, it was heat-fragmented at 95°C for 25 min and the ends were repaired with TruSeq Ribo Profile PNK (Illumina). RNA was then cleaned using a modified RNA Clean & Concentrator-5 kit (Zymo Research) method, adding 200 µl of RNA binding buffer and 450 µl of 100% ethanol before continuing with the manufacturer’s protocol.

#### Adapter ligation and library amplification

The 3’ adapters were ligated according to the TruSeq kit (Illumina) manufacturer’s instructions, followed by reverse transcription of the library. The resulting cDNA was extracted using a 10% polyacrylamide/7-8 M urea/TBE gel, circularized according to the TruSeq kit instructions, and amplified as per their protocol. The library was then cleaned with Agencourt AMPure XP beads (Beckman Coulter).

#### Gel extraction

Finally, the cDNA libraries were gel extracted using an 8% TBE gel.

#### Sequencing

Sequencing was performed on an Illumina NextSeq 500, using a NextSeq500/550 high output V2 (75 cycles) kit on SE-mode.

### Processing of chimeric eCLIP data

The raw sequencing reads of the chimeric eCLIP experiment were processed using our iCLIP/eCLIP processing pipeline `racoon_clip` (https://github.com/ZarnackGroup/racoon_clip/) (38). To achieve this, a new miR-eCLIP module was added (`racoon_clip` version 1.1.0) which allows to separate and individually process regular eCLIP and chimeric reads from the data. Regular reads are treated like a standard eCLIP dataset. In the following, the individual steps of `racoon_clip` are described in detail (**Figure S1A**). A detailed description of the usage and functions of the miR-eCLIP in `racoon_clip` and is available at: https://racoon-clip.readthedocs.io/en/latest/tutorial_mir.html.

#### Quality filtering and adapter trimming

Sequencing reads from chimeric eCLIP were filtered for a Phred score >= 10 inside the unique molecular identifier (UMI) at positions 1-10 of each read to ensure reliable sample and duplicate assignment. Then, 3’ adapters (Illumina Universal Adapter, Illumina Multiplexing Adapter, and eCLIP adapters 1-20) were trimmed with FLEXBAR (version 3.5.0) (39) using two cycles (--adapter-trim-end RIGHT--adapter-error-rate 0.1--adapter-min-overlap 1--adapter-cycles 2). At the same time, UMIs were trimmed from the 5’ end of the reads and stored in the read names (--umi-tags--barcode-trim-end LTAIL). Reads that were shorter than 15 nt after trimming were discarded (--min-read-length 15).

#### miRNA alignment

To separate regular eCLIP reads from chimeric reads generated through ligation of a miRNA and a corresponding mRNA fragment, the read starts were aligned to miRNA sequences from MiRBase (37). For this, filtered and trimmed reads were shortened to the first 24 nt with fastx_trimmer-l 24 (from FASTX-Toolkit, version 0.0.14, http://hannonlab.cshl.edu/fastx_toolkit/). The read starts were then aligned to the mature miRNA sequences from MiRBase (accessed 10.10.2022, based on the mm10 genome assembly) using bowtie2 (version 2.4.5) (40) with the following settings:--local-D 20-R 3-L 10-i S,1,0.50-k 20--trim5 2, after creating an index of the miRNA genome assembly with bowtie2-build.

The resulting alignment file was split intro read starts from chimeric and regular eCLIP reads using samtools view-f 0 (aligned) and samtools view-f 4 (unaligned), respectively (SAMtools version 1.11) (41). Then, the read names of the unaligned read starts were used to extract the complete regular eCLIP reads from the quality filtered and trimmed fastq files using seqkit grep-n (SeqKit version 2.3.1) (42) which were then sorted with seqkit sort-n.

The read starts of chimeric reads were split by the position of their miRNA mapping start (positions 1-4, column 4 in sam file) using awk (mawk 1.3.3). Then, the respective reads were extracted from the quality filtered and trimmed fastq files according to the miRNA mapping start position with seqkit grep-n. Moreover, the names of the miRNA (column 3 of sam file) were extracted using awk and added to the beginning of the read names in the fastq files using seqkit replace-p ‘(.+)’-r “{{kv}}”.

For each miRNA mapping start position, the reads in the fastq files were trimmed by miRNA mapping start plus 21 nt from the 5’ end. Thus, the trimmed reads contained the mRNA portion of the chimeric reads, such that their 5’ position corresponded to the position of the UV crosslink. After trimming, the fastq files of the chimeric reads with miRNA start position 1-4 were merged again using cat. Reads with miRNA mapping starts at later positions were discarded.

#### Genome alignment and deduplication of regular and chimeric eCLIP reads

Regular eCLIP reads and trimmed chimeric reads were aligned to the mouse genome (version GRCm38.p6) using GENCODE gene annotation version M23, CHR (43) with STAR (version 2.7.10, (44). In short, the genome was indexed with STAR –runMode genomeGenerate. Then, the regular and chimeric eCLIP reads of each sample were individually aligned to the genome with STAR –runMode alignReads (--sjdbOverhang 139 –outFilterMismatchNoverReadLmax 0.04--outFilterMismatchNmax 999--outFilterMultimapNmax 1--alignEndsType “Extend5pOfRead1”--outReadsUnmapped “Fastx”--outSJfilterReads “Unique”). Obtained bam files were indexed with samtools index. Aligned reads were deduplicated with umi_tools dedup--extract-umi-method read_id--method unique (UMI-tools version 1.1.1) (45).

#### Assignment of crosslink sites of regular and chimeric eCLIP reads

The deduplicated bam files of regular and trimmed chimeric eCLIP reads were converted into bed files using bedtools bamtobed (version 2.30.0) (46). The reads were the shifted by 1 nt upstream with bedtools shift-m 1-p-1 because the UV crosslink sites should be positioned 1 nt upstream of the eCLIP read starts. For chimeric reads, the miRNA name was extracted from the read name using awk. The bed files were split into plus and minus strand, and the reads were then reduced to 1-nt crosslink events using awk.

To allow for visualization, the bed files of 1 nt events were converted to bigWig files using bedGraphToBigWig (ucsc-bedgraphtobigwig version 377, https://github.com/ucscGenomeBrowser/kent). Additionally, the bigWig files of replicates were merged by sample type (Ago-IP WT, Ago-IP KO, miR-181-enriched WT, miR-181-enriched KO) with bigWigMerge (ucsc-bigwigmerge version 377).

#### Binding site definition

We defined binding sites using the R/Bioconductor package `BindingSiteFinder` (version 1.4.0, 10.18129/B9.bioc.BindingSiteFinder). The underlying workflow has been described in (47). In short, sites with significant pileups of crosslink events were called using PureCLIP (version 1.3.1) (48) with default parameters on the merged replicates of each sample type. The PureCLIP-called sites for Ago2-IP samples were then filtered to remove the 5% of sites with lowest PureCLIP scores (global filter), followed by a second filter to remove the 50% of sites with lowest PureCLIP score per gene (gene-wise filter). Then, binding sites were defined using `BindingSiteFinder` with bsSize=7, minWidth=2, minCrosslinks=2, minClSites=2 and filtered for reproducibility between replicates using a replicate-wise threshold at 5% that had to be met in all three replicates (**Figure S1e, f**).

We defined three sets of Ago2 binding sites (**Table S1**): (i) Ago2 binding sites in WT thymocytes were defined from regular eCLIP reads of 3 Ago2-IP WT replicates (*n* = 27,497). (ii) Ago2 binding sites in miR-181a/b-1 KO cells were defined from regular eCLIP reads of 3 Ago2-IP KO replicates (*n* = 21,852). (iii) miR-181 MREs (i.e., Ago2 binding sites after miR-181 enrichment) were defined from the union of regular eCLIP reads and mRNA parts of chimeric reads of 3 miR-181-enriched WT replicates (*n* = 6,724).

#### Assignment of miRNAs in Ago2 binding sites using chimeric reads

From the processing procedure described above, we obtained a total of 230,906 chimeric reads from the merged Ago2-IP WT replicates and 180,528 from Ago2-IP KO replicates (**Table S1**). After miR-181 enrichment, we obtained 741,279 from the merged WT replicates and 38,282 from the merged KO replicates. To assign an Ago2 binding site with a miRNA from chimeric reads, we required that the crosslink sites of the mRNA parts (i.e., 1 nt upstream of mapped mRNA fragment) of at least 1 chimeric reads with this miRNA started either inside the Ago2 binding site or within 10 nt to either side.

To assess the co-occurrence of miRNAs in Ago2 binding sites, we calculated enrichment by pairwise Fisher’s exact tests between the 10 most detected miRNAs plus miR-181c-5p and miR-181d-5p. *P* values were corrected using Benjamini-Hochberg method and clustered by the complete linkage method.

#### Assignment to genes and transcript regions

Ago2 binding sites and miR-181 MREs were assigned to genes using GENCODE gene annotation (version M23, CHR). Binding sites overlapping multiple genes of different biotypes were resolved using the hierarchy protein-coding > tRNA > lincRNA > miRNA > snRNA. Note that we use the term transcripts equivalent to genes from hereon.

Within protein-coding transcripts, the miR-181 MREs were further assigned to transcript regions, including introns, coding sequences (CDS) and 3’ and 5’ untranslated regions (UTRs). Overlaps with multiple different transcript regions were resolved by the hierarchical rule: 3’ UTR > 5’ UTR > CDS > intron. This procedure yielded the following number of MREs per region: 3’ UTR – 2,343, 5’ UTR – 379, and CDS – 1,936. 1,765 intronic sites were excluded from further characterization. Density distribution of MRE sites along gene regions were plotted with the cliProfiler package (version 3.6.1, doi:10.18129/B9.bioc.cliProfiler).

#### Differential Ago2 binding between WT and KO cells

For the differential binding analysis, Ago2 binding sites from the Ctrl and the KO condition were combined. There were three possible scenarios: (i) Binding sites that were identical in both datasets, i.e., having the same start and end coordinates, were directly transferred to the merged binding site set. (ii) Binding sites that did only occur in one dataset were also directly included in the merged binding site set. (iii) If binding sites occurred in both datasets but were slightly shifted towards each other, i.e., having overlapping but not identical coordinates, these were recentered into one concordant binding site centred on the position with the highest number of crosslink events in the merged signal from both conditions. This yielded a combined set of 28,253 Ago2 binding sites, which had been found in either WT, KO or both conditions.

To test for differential Ago2 binding, we used DEseq2 (version 1.38.3) (49) with a two-factor design containing both the number of Ago2 crosslink events within each Ago2 binding site (labeled “type”) and the background crosslink events in the remainder of the transcript (labeled “condition”). For counting background crosslink events, an offset of 5 nt to either side of the Ago2 binding sites was excluded. To exclude transcripts with low overall signal, we calculated expression estimates (TPM-like) based on the number of crosslink events in the background and removed all transcripts that were not expressed in both conditions using a TPM cutoff > 50, thereby keeping 26,462 merged Ago2 binding sites in 3,779 transcripts for subsequent analysis (**Table S3**). The analysis was performed using the AGO2-IP samples from miR-181 KO vs. WT conditions (3 replicates each).

We tested for significant log_2_ fold changes in Ago2 binding (crosslink events) using the likelihood ratio test (LRT) of DESeq2 with the design formula “design = ∼condition + type + condition:type”, where the interaction term “condition:type” represented log_2_ fold changes of binding sites that were due to transcript-level changes between the conditions. As a result, the log_2_ fold changes per binding site were corrected for a potential change in transcript expression. *P* values were adjusted by independent hypothesis weighting using the R/Bioconductor package IHW, version 1.26.0) (50). In total, we identified 1,074 significantly changed Ago2 binding sites (adjusted *P* value < 0.05), including 278 (25.9%) upregulated and 796 (74.1%) downregulated sites. These included 1,055 sites on protein-coding transcripts that were distributed across transcript regions as follows: 3’ UTR – 391 (106 up / 285 down), 5’ UTR – 63 (32 up / 31 down), CDS – 268 (75 up / 193 down), intron – 303 (53 up / 253 down) (**Figure S1j, Table S2**).

### Gene Ontology (GO) analysis

GO enrichment analysis was performed using the enrichGO function in the cluster profiler package (version 4.8.2) (51) and then reducing the number of categories with REVIGO (52), using the parameter settings “small (0.5)” and “SimRel”. The resulting list was further manually curated for redundancies and terms not applicable to lymphocytes. The full lists can be found in **Table S4**.

### Differential transcript abundance in RNAseq and RF

Total RNA reads were checked for quality using FastqC and adapters were removed using Flexbar (version 3.5.0, options:--min-read-length 15-ao 1, adapter sequence: “AGATCGGAAGAGCACACGTCT”) (53). Reads corresponding to rRNA were removed with Bowtie2 (version 2.5.1, options:--very-sensitive-x rRNA) (40) and remaining reads were aligned to the mouse genome (GENCODE M23) using STAR (version: 2.7.10a, options:--outFilterMultimapNmax 5--alignEndsType EndToEnd) (44). Reads per gene were counted using htseq-count (version 2.0.3) (54) with default settings (--stranded=yes) and GENCODE gene annotation (version M23). Batch effects were removed using the ComBat-seq function of the sva package (version 3.48.0) (55). Differential expression analysis between miR-181a/b-1 KO and WT samples was performed using DESeq2 (version 1.40.2) (50) with a significance cutoff at an adjusted *P* value < 0.05 (Benjamini–Hochberg correction). This yielded a total of 548 significantly differentially abundant transcripts in RNAseq, including 327 with increased and 221 with decreased expression upon miR-181 KO, and 1,207 significantly differentially abundant transcripts in the ribosome footprint, with 711 increased and 496 decreased. Statistical testing for ECDF plots was performed using the Kolmogorow-Smirnow test.

### Translational efficiency (TE)

Translational efficiency was calculated using deltaTE as outlined in paper (56), using DEseq2 (version 1.40.2) (50). Welch’s two-sample *t*-test was used to test for significance in density plots.

### Motif analyses

All motif and RNA structure analyses were performed using mature transcript sequences, and miR-181 MREs were transferred to transcript coordinates.

#### Assignment to mature transcript isoform

miR-181 MREs were assigned to a specific transcript isoform by the following criteria: First, we selected all transcript isoforms which contained the MRE in the assigned transcript region (see above). Intronic MREs were excluded from this analysis and mature transcripts were used. If this yielded more than one transcript isoform, protein-coding transcripts were preferred and the transcripts with the highest transcript support level were chosen. If more than one transcript remained, the longest transcript was selected. miR-181 MREs were mapped to the relative coordinates on the respective transcript transcripts using mapToTranscripts() with the parameter extractor.fun=GenomicFeatures::exonsBy from the R/Bioconductor GenomicFeatures (version 1.50.4) (57). Using this procedure, 4,519 miR-181 MREs could be mapped to a mature transcript.

#### Obtaining fasta of regions of interest

Sequences of mature transcripts are obtained from GENCODE (“transcript sequences” fasta file) provided together with the genome assembly (GRCm38.p6). These sequences were used as input for the getSeq function of the R/Bioconductor package Biostrings (version 2.66.0, DOI: 10.18129/B9.bioc.Biostrings) which extracts sequences in the windows of interest. Fasta files of the sequences of interest were then written with to writeXString() function.

#### De novo motif discovery

We used XSTREME to discover motifs *de novo* in a 401 nt window surrounding of the miR-181 MREs considering mature transcripts. The 401 nt window was chosen on an empirical basis with other window sizes between 51 and 1001 nt showing very comparable results (**Figure S8**). XSTREME was executed via the MEME SUITE webpage (https://meme-suite.org/meme/tools/xstreme, MEME version 5.5.3, Sat Jun 10 16:19:38 2023-0700) (58). The parameters for this task are as follows: E-value must be less than or equal to 0.05, width should be between 5 and 10, and the background consists of model control sequences. The STREME limit was set to 20 motifs, MEME options were set to a “Default E-value” and “Zero or one occurrence per sequence”. The motif with the lowest E-value is shown.

#### Counting occurrences of motif variations

We searched for variations of the seed motif in a 200-nt window downstream of the start position of the miR-181 MREs with the vmatchPattern() function from BioStrings (8mer: “UGAAUGUA”, 7mer_m8: “UGAAUGU”, 7mer_a1: “GAAUGUA”, 6mer: “GAAUGU”, 8mer with wobble position: “UGAUUGUA”, 7mer_m8 with wobble position: “UGAUUGU”, 7mer_a1 with wobble position: “GAUUGUA”, 6mer with wobble position: “GAUUGU”, alternative 6mer (3–8): “UGAAUG”, alternative 6mer (3–8) with wobble position: “UGAUUG”). The window size was established empirically based on the crosslink distribution relative to the binding sites (**Figure S8**). The start position of all 6mer motifs relative to the MRE is shown.

We then selected for each miR-181 MRE the seed match with the least distance to the binding site. The first seed per miR-181 MRE was used to count the occurrence of seed match types.

### Analysis of satellite repeat elements

Repeat sequences were obtained from RepeatMasker (Smit, A., Hubley, R. and Green, P., RepeatMasker Open-4.0, http://www.repeatmasker.org) by selecting the annotation “AH99012” from the R/Bioconductor package AnnotationHub (version 3.6.0, doi:10.18129/B9.bioc.AnnotationHub). The RepeatMasker annotation was the lifted to mature transcript coordinates as described above and overlapped with the miR-181 MREs.

### Hybrid structures of miR-181 with its target RNAs

We predicted hybrid structures of miR-181a to the miR-181 MREs with RNAduplex (ViennaRNA Package version 2.4.17) (59). In short, RNAduplex was run with default parameters using the mature miR-181a sequence together with the transcript sequence from 30 nt before until 200 nt after each miR-181 MRE. For the MRE transcript coordinates and obtaining sequences in fasta format, see Motif analyses.

Sequences were split into MREs with or without an AA seed match in the considered window (**Figure S9a**). After duplex prediction, sequences with a seed match were split into those in which the predicted duplex did or did not occur within the seed match. The former were considered as canonical seed pairing (408 MREs). Sequences without a genuine seed match were split according to whether the duplex did or did not involve partial pairing within the first 8 nt (**Figure S9d**). The minimum free energy was normalized by dividing by number of miR-181a nucleotides included in the duplex structure.

*k*-means clustering was performed on the dot-bracket annotation of the predicted miR-181a duplex structures. For this “.” was turned to 0 and “)” was turned to 1. To prioritize sequential binding without gaps, sequential 1’s without intervening 0’s were added up. For example, “.))).” would be turned to “03330”. *k*=8 was empirically chosen in order to optimize the visualization.

## Results

### Chimeric eCLIP identifies Ago2 binding sites in murine thymocytes

To determine the miR-181a/b targetome in murine thymocytes, we captured trimeric Ago2– miRNA–mRNA interactions in a two-step procedure termed chimeric enhanced UV crosslinking and immunoprecipitation (chimeric eCLIP) (60) (**Figure 1a**). In the first step, Ago2–RNA complexes were covalently crosslinked by UV irradiation and then immunoprecipitated with an Ago2-specific antibody (referred to as Ago2-IP). To preserve the miRNA–mRNA pairing, the miRNA and mRNA fragments in the complexes were then ligated into chimeras. In the subsequent miRNA enrichment, miR-181a/b-containing chimeras were selectively enriched using antisense oligonucleotides (referred to as miR-181-enriched). High-throughput sequencing of the Ago2-IP libraries yielded a mixture of regular eCLIP reads and chimeric reads, which informed on all Ago2 target sites and the associated miRNAs, respectively, while the miRNA-enrichment samples were used to specifically identify the miR-181 MREs (see below). Experiments were performed on total murine thymocytes isolated from wildtype (WT) and miR-181a/b-1 knockout (KO) mice (29) (3 biological replicates), yielding an average of ∼50 million (mio) reads per sample (**Table S1**).

**Figure 1.**
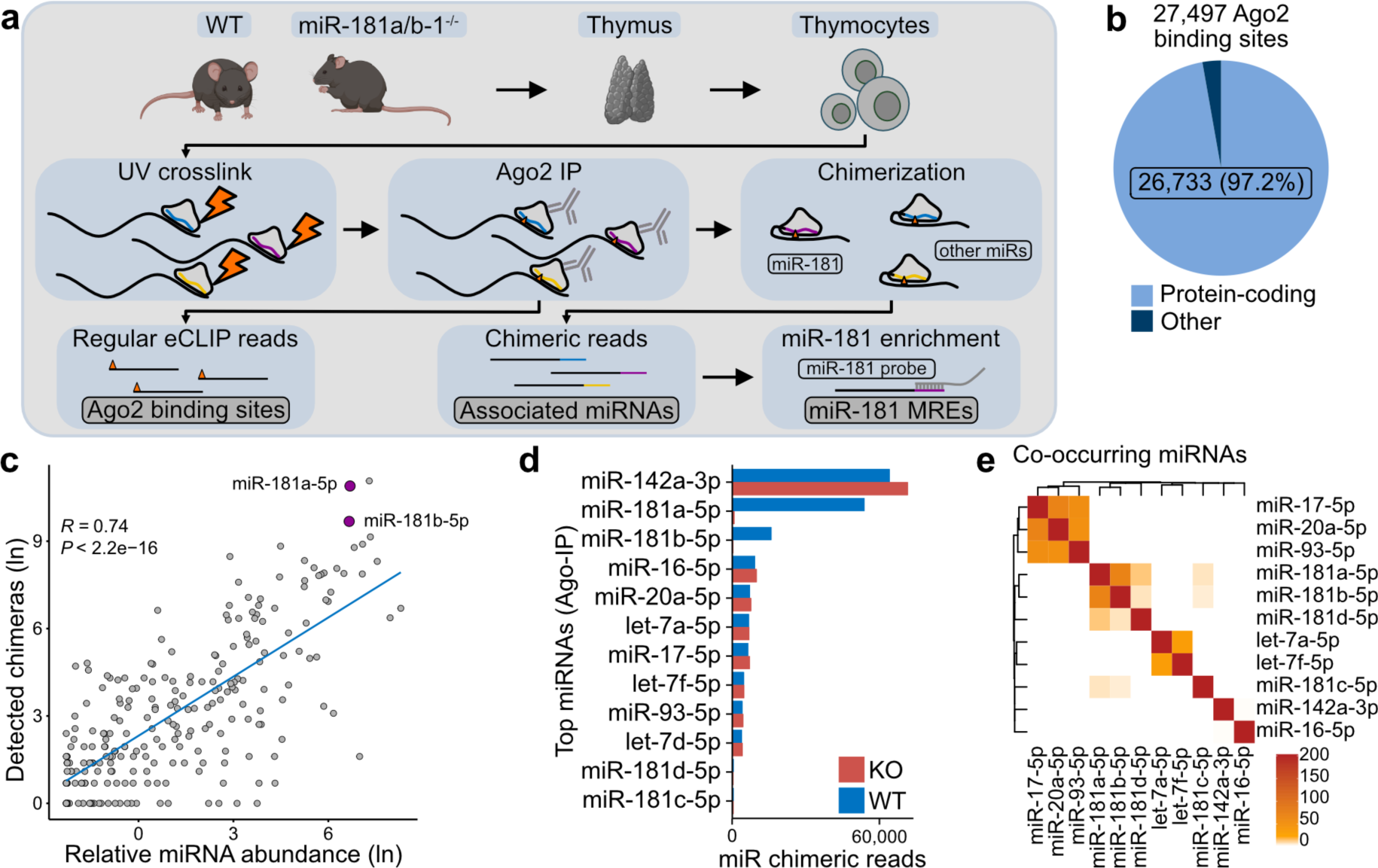
Chimeric eCLIP identifies Ago2 binding sites in murine thymocytes. **(a)** Chimeric eCLIP experiment to map Ago2 binding sites and miR-181 MREs. Top: Thymocytes were extracted from thymi of wild-type (WT) and miR-181a/b-1 knockout (KO) mice. Middle: In the extracted cells, Ago2 is crosslinked to its target RNAs by UV irradiation, followed by Ago2 immunoprecipitation (IP). Chimerization ligates miRNAs and mRNA fragments in the Ago2-RNA complexes before reverse transcription. Bottom: This results in two types of reads: regular eCLIP reads that truncate at the Ago2 crosslink sites (right) and chimeric reads that consist of the miRNA followed by the target RNA sequence (left). Chimeric reads harboring miR-181 are selectively enriched using a complementary probe to define miR-181 MREs. **(b)** Distribution of Ago2 binding sites (*n* = 27,497) in protein-coding (lightblue) and other (darkblue) transcript biotypes. **(c)** Comparison of miRNA counts in chimeric reads in Ago2-IP from WT cells with abundance in thymus cells from PCR measurements (26). miR-181a-5p and miR-181b-5p are highlighted (purple). Pearson correlation coefficient and associated *P* value are given. Blue line displays smoothed conditional means using loess. **(d)** Number of chimeric reads for the 10 most abundant miRNAs in the Ago2-IP samples from WT (blue) and KO (red) cells. miR-181c-5p and miR-181d-5p were added for comparison. **(e)** Co-occurrence of miRNAs in Ago2 binding sites. Pairwise enrichment between miRNAs in (e) was calculated by Fisher’s exact test. Color scale gives adjusted *P* value (Benjamini-Hochberg) on negative logarithmic scale.

To efficiently process the chimeric eCLIP data, we made use of our automated iCLIP/eCLIP processing pipeline `racoon_clip`) (**Figure S1a**) (38). This pipeline includes all steps from quality control, demultiplexing and barcode handling, genome alignment, deduplication, and assignment of single-nucleotide crosslink sites. We added a new module for chimeric eCLIP to specifically handle chimeric reads, assign a miRNA annotation, and consistently identify crosslink events for both chimeric and regular eCLIP reads. All steps can be customized to best fit the data at hand.

We first used the data from the initial Ago2 IP to globally assess RISC binding and miRNA loading for all expressed miRNAs. To this end, we used the regular eCLIP reads from both WT and KO samples to identify Ago2 binding sites using the R/Bioconductor package `BindingSiteFinder` (47) (**Figure S1b**). Rather than taking complete read pileups (60), we narrowed down the Ago2 binding sites to an optimal width of 7 nt (**Figure S1c, d**). In total, we identified 27,497 reproducible Ago2 binding sites (**Figure S1e, f**). The vast majority of these (26,733, 97%) were located in the transcripts of 4,589 protein-coding genes (**Figure 1b**).

Next, we used the chimeric reads from the Ago2-IP samples to investigate which miRNAs were present in the Ago2–mRNA complexes. In the Ago2-IP samples from WT mice (Ago2-IP WT), we found a total of 230,906 uniquely mapping chimeric reads after PCR duplicate removal, corresponding to 317 different miRNAs. Their frequency correlated well with published miRNA expression levels in double-positive thymocytes (26) (**Figure 1c**), indicating that chimeric eCLIP quantitatively captures the miRNA loading of RISCs in cells. Particularly, miR-142-3p (28%) and miR-181a-5p (23%) together accounted for more than half of all detected miRNAs (**Figure 1d**). In total, 26% (7,052) of all Ago2 binding sites could be associated with a specific miRNA. While most Ago2 binding sites harbored just one miRNA, 8% contained two or more distinct miRNAs (**Figure S1g**). The commonly co-occurring miRNAs shared the same seed sequence (**Figure 1e**), suggesting that RISCs loaded with miRNA family members frequently compete for the same MREs. In the chimeric reads from KO cells (Ago2-IP KO), miR-181a-5p and miR-181b-5p were selectively lost, while miR-142-3p and all other miRNAs remained stable (**Figure 1d**). Consistent with earlier studies, these data indicate that miR-181a-1 and b-1 are much more abundantly expressed than their miR-181a-2 and b-2 counterparts (29). Chimeras containing miR-181c and miR-181d as well as the 3p strands of any miR-181 family member were low to undetectable, which is consistent with low levels of expression and documented strand dominance (26,37). Hence, based on these results and the chosen experimental approach, MREs characterized in this study mostly reflect binding sites for miR-181a-1-5p and miR-181b-1-5p. For reasons of clarity, we refer to those as miR-181 MREs, unless specifically stated otherwise. Altogether, these results supported that Ago2 universally targets mRNAs in murine thymocytes and that miRNA abundance predominantly drives the extent of RISC binding. They confirmed that miR-181 is a predominant miRNA in these cells and indicated that loss of miR-181a/b-1 does not result in substantial compensatory takeover by selective other miRNAs.

### miR-181 targets hundreds of MREs in the transcriptome

To comprehensively identify the MREs targeted by miR-181, we used the miR-181-enriched samples, in which miR-181a/b-containing chimeras had been pulled down using antisense oligonucleotides (**Figure 1a**). In line with a high enrichment efficiency, the 741,279 chimeric reads from the WT samples contained almost exclusively miR-181a (80%) and miR-181b (18%; **Figure 2a, Table S1**). In contrast, the KO samples yielded only 38,282 chimeric reads which harbored residual background sequences. Using the WT samples, we identified 6,724 reproducible miR-181 MREs in 2,995 target transcripts (**Figure S1h, i**). Of note, 80% of the miR-181 MREs had not been detected in the initial Ago2-IP (**Figure 2b**), underlining that the oligonucleotide-based enrichment massively increased the number of detected MREs.

**Figure 2.**
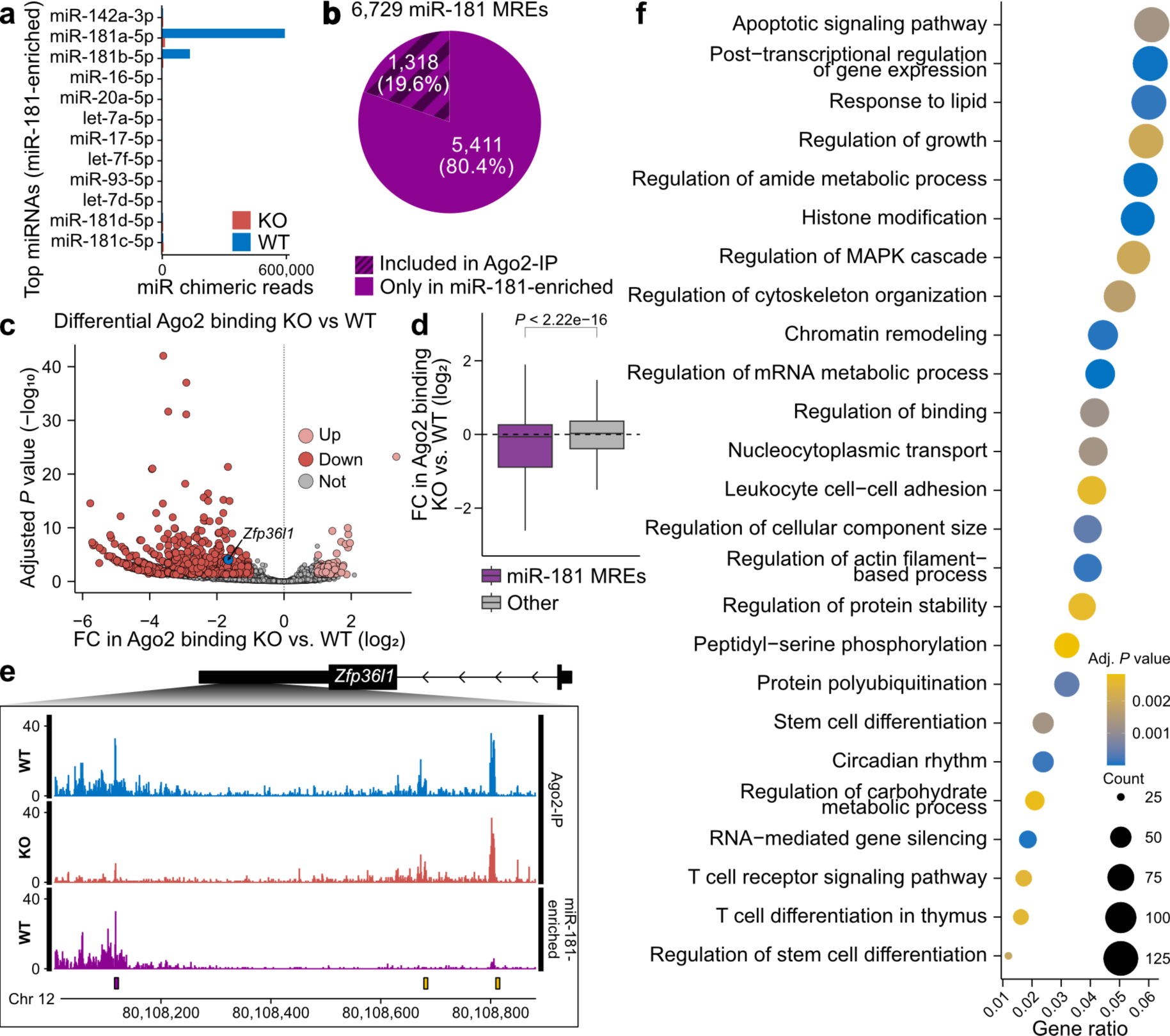
miR-181 targets hundreds of MREs in the transcriptome. **(a)** Number of chimeric reads for the 10 most abundant miRNAs for miR-181-enriched samples from WT (blue) and KO (red) cells. miR-181c-5p and miR-181d-5p were added for comparison. **(b)** miR-181 MREs from miR-181-enriched WT samples (*n* = 6,729). Of these, 1,318 MREs (19.6%, dark shading) had also been detected by miR-181-containing chimeric reads in Ago2-IP WT samples. **(c)** Differential Ago2 binding in WT and KO cells. Volcano plot shows log2-transformed fold change (FC) against adjusted *P* value (independent hypothesis weighting, IHW). Significantly changed Ago2 binding sites (*P* value < 0.05) are colored in red (down) or salmon (up), and all others in grey. Ago2 binding site in *Zfp36l1* shown in (b) is highlighted in blue. **(d)** Differential Ago2 binding at miR-181 MREs (purple) and other Ago2 binding sites (grey). *P* value from Welch two sample t-test shown above. **(e)** Genome browser view of chimeric eCLIP data in the 3’ UTR of *Zfp36l1*. Shown are crosslink events (merged replicates) of Ago2-IP from WT (blue) and KO (red) cells as well as after miR-181 enrichment from WT cells (purple). Identified Ago2 binding sites are show below, colored by miR-181 MRE (purple) and other (yellow). **(f)** GO analysis of transcripts containing miR-181 MREs. GO terms were semantically reduced using REVIGO followed by manual curation. The top 25 GO terms ranked according to adjusted *P* values are shown, with gene count in each set visualized by dot size and adjusted *P* values by colour gradient.

To confirm that the identified MREs were indeed targeted by miR-181, we tested for changes in Ago2 binding in the initial Ago2-IP samples from miR-181a/b-1 KO cells. Using a differential binding analysis that corrects for gene expression changes, we detected a significant downregulation in 796 Ago2 binding sites (74% of all changing Ago2 binding sites; **Figure 2c, Figure S1j, Table S2**). Moreover, Ago2 binding globally went down at the miR-181 MREs, while all other Ago2 binding sites showed no such shift (**Figure 2d**). The reduced Ago2 binding in the KO cells strongly supported the notion that RISC targeting to the identified miR-181 MREs is indeed dependent on miR-181 presence.

Target prediction tools and compilations of experimentally suggested miRNA targets, such as TargetScan and miRTarBase, respectively, list hundreds of potential targets for miR-181a/b (61,35). Out of a total of 2,995 transcripts containing miR-181 MREs in our study, 362 and 102 are listed exclusively in TargetScan and miRTarBase, respectively (**Figure S2a**). 86 mRNAs are included in both databases. The overlap with each database correlates with the total number of targets therein, i.e., a similar percentage of the miR-181a/b targets present in TargetScan (40%) and miRTarBase (46%) harbored miR-181 MREs from our study. Moreover, we note that the agreement between both databases is generally low, as only 187 putative target mRNAs are listed in both miRTarBase and TargetScan. To further evaluate the overlap with TargetScan, we compared mean expression levels of miR-181a/b target mRNAs in our RNAseq data from WT thymocytes in which the eCLIP experiment had been performed. TargetScan targets that had not been detected in our eCLIP data displayed significantly lower expression levels than eCLIP-detected targets or targets listed in both (**Figure S2b**). This suggests that the absence of targets predicted by TargetScan is due to low abundance of these mRNAs in thymocytes.

Among the MRE-containing transcripts were multiple known targets of miR-181 in thymocytes, including *Bcl2*, *Cd69*, transcripts of DUSP family members, *Ptpn22*, *Pten* and *S1pr1* (24,31,30,32) (**Table S3**). In addition, we detected MREs in the mRNAs of proteins with defined roles in T-cell development, such as Cyclin D3 or the RNA-binding proteins Zfp36l1 and Zfp36l2 (**Figure 2e**) (62–64). Consistent with earlier studies, gene ontology (GO) analysis identified enriched functionalities associated with T-cell receptor signaling, MAPK signaling and immune system development as well as apoptosis (**Figure 2f, Table S4**) (24,31,32). We also noted an enrichment of transcripts associated with intracellular processes, including representation of GO terms associated with chromatin remodeling, RNA processing as well as RNA-dependent post-transcriptional gene regulation, suggesting that miR-181 may play a role in gene-regulatory processes at a more global scale.

Thus, by combining selective miR-181a enrichment with miR-181a/b-1 deficiency, we were able to faithfully identify 6,729 miR-181 MREs in 2,995 target transcripts. Of note, this exceeds the number of *in silico* predicted targets via TargetScan by almost 3-fold (61). The expanded miR-181 targetome confirms its well-known impact on negative regulators of T-cell receptor signaling and survival factors, and newly links miR-181 to global gene regulatory processes at the level of DNA and RNA.

### miR-181 acts primarily through transcript destabilization

miRNA-mediated gene regulation can act at the level of RNA stability, translational inhibition or both. To address the primary effect of miR-181 in thymocytes, we combined RNA sequencing (RNAseq) and ribosome footprinting (RF) to capture miR-181-dependent changes in the transcriptome and translatome in WT and KO cells (**Figure 3a, Figure S3**). In RF, ribosomes are fixed on mRNAs followed by digestion of unprotected RNA. High-throughput sequencing of the ribosome-protected fragments thus allows the identification of mRNAs in the process of translation, whereby ribosome occupancy serves as a quantitative measure of translation.

**Figure 3.**
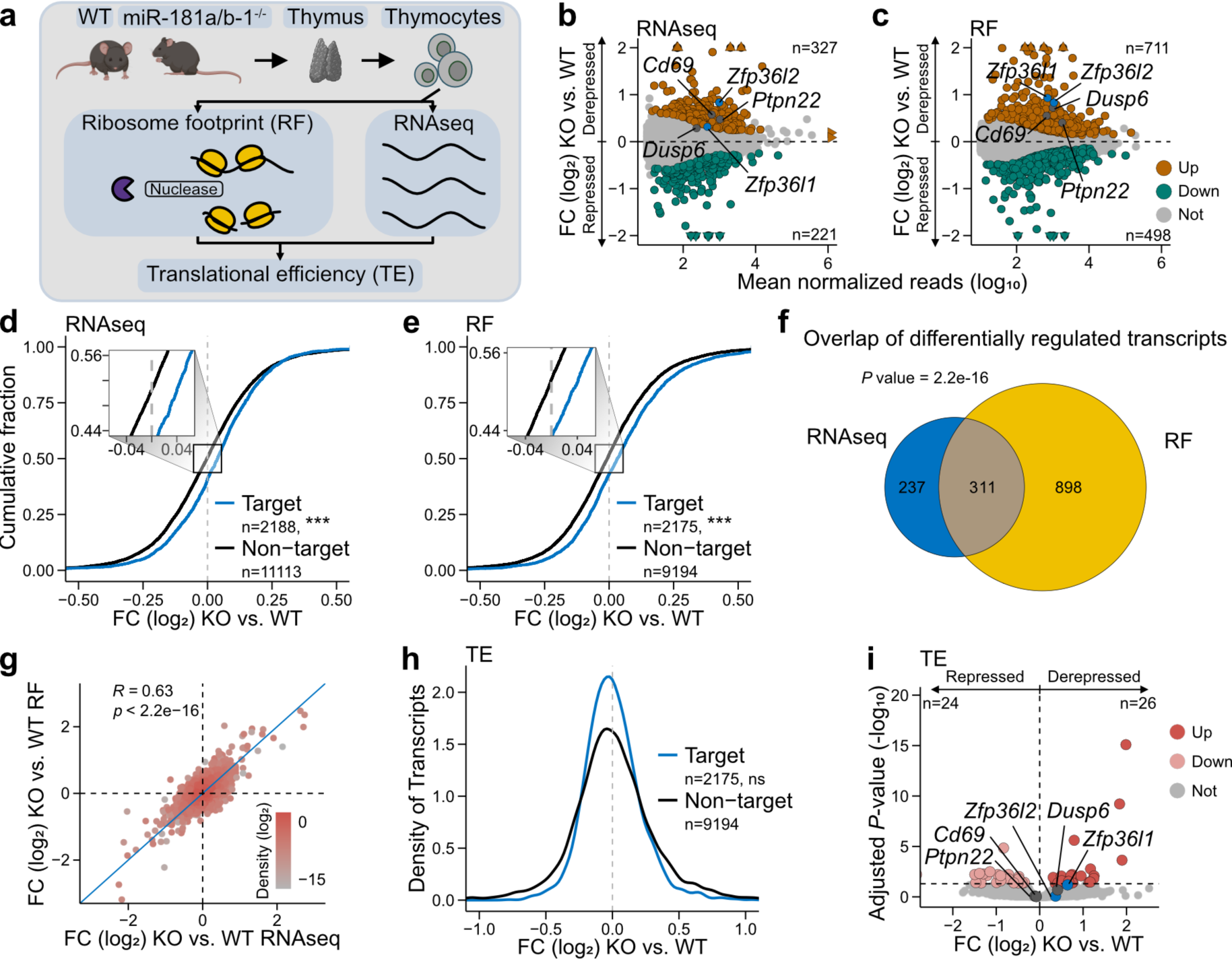
miR-181 acts primarily through transcript destabilization. **(a)** Scheme of transcriptome and translatome determination in primary thymocytes from miR-181 KO and WT mice by combined ribosome footprinting (RF) and RNAseq. RF and RNAseq data are used to determine translational efficiency (TE). **(b,c)** Transcripts with significant differences (*P* value < 0.05, Benjamini-Hochberg correction) in RNA abundance (RNAseq) (b) or ribosome occupancy (RF) (c) are plotted as log_2_ fold change (FC) of KO over WT vs. mean normalized read count. Example targets are highlighted in blue. **(d,e)** Fold changes in transcript levels (e) and ribosome occupancy (f) between miR-181 KO and WT conditions are shown as the cumulative density of the log_2_ values, for transcripts with MREs vs. non-targets. Differences between curves were tested for significance using the asymptotic two-sample Kolmogorov-Smirnov test (ns>0.05, *<0.05, **<0.01, ***<0.001). **(f)** Overlap of mRNAs with differentially abundance (RNAseq, blue) and differential ribosome occupancy (RF, yellow). *P* value from Fisher’s exact test. **(g)** Correlation of log_2_ fold-changes of KO over WT of ribosome occupancy (RF) vs. transcript levels (RNAseq). **(h)** Density plot shows changes in translational efficiency (TE) for transcripts with MREs vs. non-targets shown as log_2_ fold-changes of KO over WT. Differences between curves were tested for significance using Welch two-sample *t*-test (ns>0.05, *<0.05, **<0.01, ***<0.001). **(i)** Transcripts with differential TE in KO vs. WT. Volcano plot shows log_2_-transformed fold change (FC) against adjusted *P* value (IHW).

In total, we detected 548 and 1,209 differentially regulated transcripts in the RNAseq and RF data, respectively (**Figure 3b, c, Table S5**). In both datasets, a larger proportion of differentially regulated transcripts was upregulated in the KO thymocytes, (**Figure 3b, c**). These comprised previously characterized targets, such as *Cd69*, *Ptpn22*, *Dusp6*, as well as the newly identified targets *Zfp36l1* and *Zfp36l2*. Effect sizes were mostly low, in line with the principally small effects on individual mRNAs as reported previously (65). The downregulated transcripts as well as a certain proportion of upregulated transcripts were likely to reflect indirect effects, including the altered composition of thymocyte populations in miR-181a/b-1 KO mice. Importantly, the miR-181 MRE-containing transcripts (referred to as targets) showed a global shift towards upregulation in both RNAseq and RF, supporting that the identified miR-181 MREs are indeed functional (**Figure 3d, e**).

A direct comparison between transcriptome and translatome showed a significant overlap of differential transcripts and a strong positive correlation in both datasets (**Figure 3f, g**). The concordant behavior indicated that the observed changes were primarily driven by differences in transcript abundance rather than translation. To test this, we used the algorithm deltaTE to calculate translational efficiency for each mRNA based on the ratio of RNAseq and RF fold changes (56). Indeed, the global translational efficiency of miR-181 targets broadly overlapped with that of non-targets, and hardly any transcripts showed a significant change in translational efficiency (**Figure 3h, i**). This suggested that miR-181 modulates gene expression in thymocytes predominantly at the level of transcript destabilization rather than translational repression.

Next, we assessed the functionality of novel vs. previously annotated and predicted targets in miRTarBase and TargetScan, respectively. In KO thymocytes, mRNAs detected by eCLIP and predicted by TargetScan or listed in miRTarBase displayed stronger derepression than mRNAs from either group alone in both RNAseq and ribosome footprinting (**Figure S2c, d**). Nevertheless, targets identified through eCLIP alone still displayed a certain degree of derepression. Expectedly, targets not predicted by TargetScan were enriched for the absence of seed matches as well as the presence of the alternative seed match identified here. However, this fraction of mRNAs also contained a large proportion of seed-match-positive mRNAs and those containing multiple MREs (**Figure S2e**). Hence, we conclude that the eCLIP-identified targets expand the pool of functional miR-181a/b targets. The combination of physical binding shown by chimeric eCLIP and orthogonal information from prediction algorithms or prior biological evidence further increases the power to detect such targets.

Taken together, we observed a substantial upregulation of transcripts in the transcriptome and translatome in response to miR-181a/b-1 deficiency, supporting a global repressive function of miR-181 in thymocytes. The detected changes clearly depended on the presence of miR-181 MREs, underlining that the identified miR-181 MREs are functionally relevant. Integration of transcriptome and translatome strongly supports the conclusion that miR-181 primarily controls mRNA abundance rather than translational efficiency.

### Number and location of miR-181 MREs determine functionality

In order to determine the functional parameters of miR-181 MREs (hereafter referred to as MREs), we focussed on MREs in mature mRNAs (**Figure 4a**). Whereas the majority contained a single MRE (1,159; 53%), 491 mRNAs (23%) contained two MREs and a similar number contained three or more MREs (525; 24%; **Figure 4b**). Using the translatome data as a functional readout, we found that a single MRE was sufficient to confer repression. More MREs progressively increased the effect (**Figure 4c, Figure S4a, b**). To validate the dose-dependent regulation by miR-181, we employed a dual reporter system driving expression of a control fluorescent protein (mCherry) and GFP from a bi-directional promoter in Scid.adh.2c2 cells ectopically expressing miR-181a (66,67). The GFP open reading frame was fused to a synthetic 3’UTR encoding 0–3 8mer miR-181-5p seed matches in various combinations (**Figure S5a**). Reporter assays showed that increasing the number of MREs resulted in progressively lower GFP levels, underlining the dose dependency of miR-181a/b-5p-mediated repression (**Figure S5b**).

**Figure 4.**
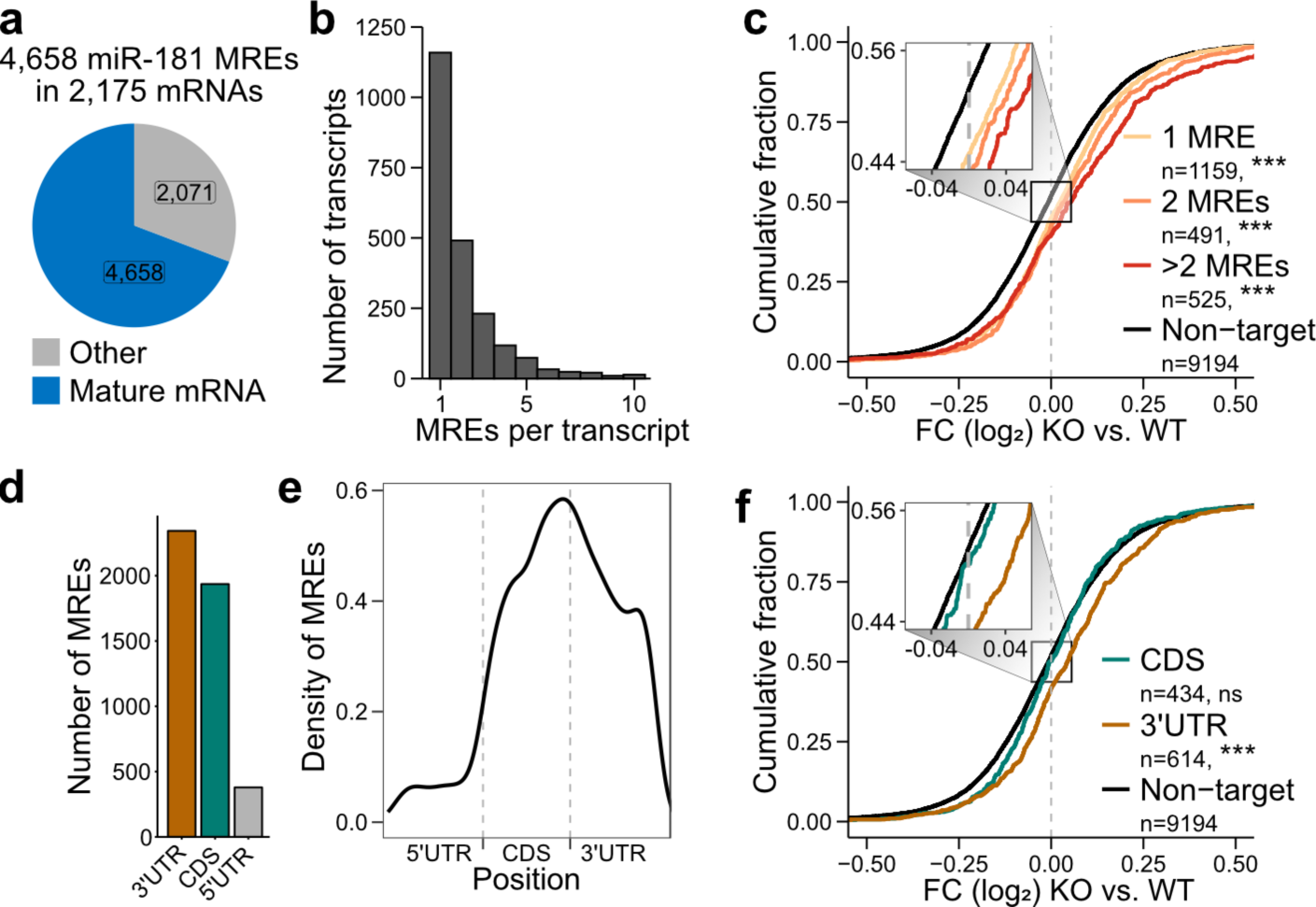
Number and location of miR-181 MREs determine functionality. **(a)** Number of miR-181 MREs located on mature mRNAs (blue) or other RNAs (grey). **(b)** Number of genes (y-axis) containing a given number of miR-181 MREs per gene (x-axis). **(c)** Fold changes in ribosome occupancy between miR-181 KO and WT conditions are shown as the cumulative density of the log_2_ values, for transcripts with one (yellow), two (orange), more than two (red) or no miR-181 MREs (black). Box on the left depicts zoom-in. Differences between curves were tested for significance using the asymptotic two-sample Kolmogorov-Smirnov test (ns>0.05, *<0.05, **<0.01, ***<0.001). **(d)** Number of miR-181 MREs per gene region. **(e)** Density of miR-181 MREs across the gene regions. **(f)** Fold changes in ribosome occupancy between miR-181 KO and WT conditions are shown as the cumulative density of the log_2_ fold changes, for transcripts with MREs in the CDS (yellow), 3’UTR (blue), or without MREs (black). Numbers of transcripts in each set are given. Box depicts zoom-in. Only transcripts with a single MRE were used. Differences between curves were tested for significance using the asymptotic two-sample Kolmogorov-Smirnov test (ns>0.05, *<0.05, **<0.01, ***<0.001).

miRNAs are expected to target mRNAs mainly via their 3’UTR (7), and commonly used algorithms such as TargetScan exclusively predict MREs in this region (61). However, we detected an almost equally large fraction of MREs in the coding sequence (CDS, **Figure 4d**) which harbored 1,936 MREs (42%) compared to 2,343 (50%) in the 3’UTR, while only a few MREs mapped to 5’UTRs (379; 8%). Within the CDS, the MRE density increased towards the stop codon (**Figure 4e**). In order to compare the potency of MREs dependent on localization, we stratified transcripts containing one MRE in either, 5’UTR, CDS or 3’UTR. Integration with the functional readout showed that, consistent with canonical models of miRNA-mediated regulation, MREs in the 3’UTR had a strong repressive capacity (**Figure 4f, Figure S4c, d**). Although we noted a subtle effect of single MREs in the CDS, this was not statistically significant.

### miR-181 MREs in satellite repeats mediate translational inhibition of zinc finger genes

To reconcile the presence of a large number of MREs with the observed limited efficacy in gene regulation, we took a closer look at the CDS MREs. We noted that some MREs accumulated in repetitive sequences from the satellite repeat families MMSAT4 and MurSatRep1 (**Figure 5a, c**). Of note, MMSAT4 repeats have been linked to the massive expansion of genes encoding KRAB-containing poly-zinc finger proteins (KZFPs), which form the largest family of transcription factors in mammalian genomes (68). Intriguingly, we found that the MMSAT4 sequence contained a site with complementarity to the miR-181 seed sequence (seed match; **Figure S6a**), which had thereby been introduced into the evolving KZFP genes. The same apparently occurred via the second type of satellite repeats, MurSatRep1, which also harbored a miR-181 seed match (**Figure S6a**) and frequently coincided with the MMSAT4 repeats (**Figure 5b**). Consistent with MREs in both repeat types were preferentially localized in the CDS, but displayed distinct positioning, with MMSAT4 MREs shifted towards the stop codon (**Figure 5d**).

**Figure 5.**
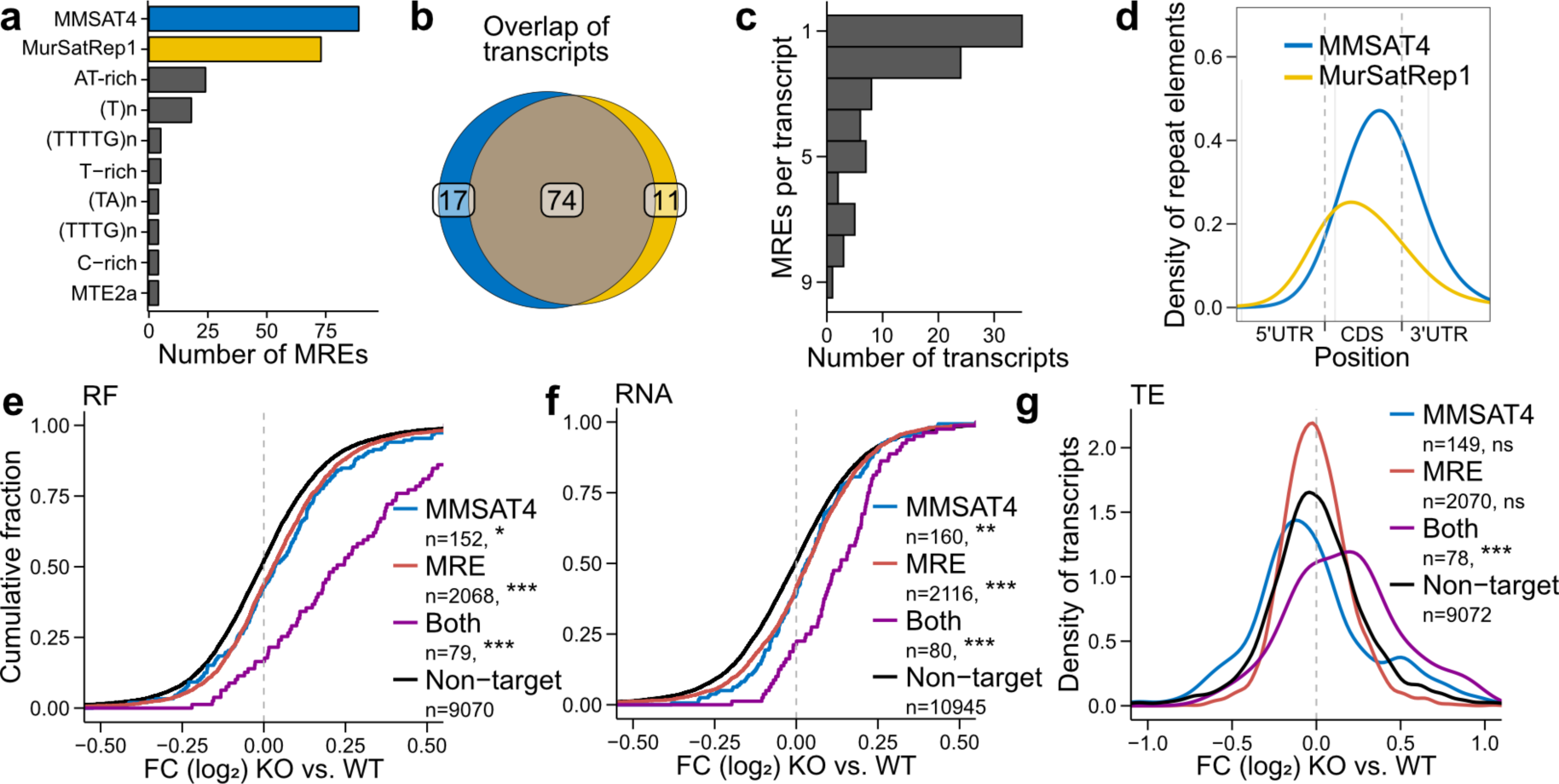
miR-181 MREs in satellite repeats mediate translational inhibition of zinc finger genes. **(a)** Left: Number of MREs overlapping with repeat elements. Right: Repeat element annotation from RepeatMasker. Top 10 overlapping repeat elements are shown. **(b)** Overlap of transcripts with Ago2-bound MREs in MMSAT4 repeat regions (blue) and MurSatRep1 regions (yellow). **(c)** Number of transcripts is counted by the number of MREs they contain. **(d)** Density of MMSAT4 (blue) and MurSatRep1 (yellow) MREs across gene regions. **(e–g)** Fold changes in ribosome occupancy (e) and RNA expression (f) are shown as cumulative density of the log_2_ fold changes for transcripts with MMSAT4 repeats (blue), transcripts with MREs (red), transcripts with both MREs and MMSAT4 repeats (purple), and non-targets (black). Differences between curves were tested for significance using the asymptotic two-sample Kolmogorov-Smirnov test (ns>0.05, *<0.05, **<0.01, ***<0.001). Translational efficiency (TE) (g) is shown as density. Differences between curves were tested for significance using the Welch two sample t-test (ns>0.05, *<0.05, **<0.01, ***<0.001).

Functional analysis revealed that the repeat-derived MREs had a massive impact on gene expression, since more than 80% of transcripts with MMSAT4 MREs were upregulated upon miR-181 KO (**Figure 5e, f**). Even in cases we had not detected Ago2 binding on the transcript in chimeric eCLIP, the presence of an MMSAT4 repeat alone was sufficient to confer regulation, with effect sizes similar those of all miR-181 MREs in other locations, suggesting that, despite the substantial depth, at least some functional MREs have eluded our analysis. The same was observed for the MurSatRep1-derived MREs (**Figure S6b, c**). Although the effect of both repeats cannot be discriminated due to the co-occurrence, the repeat elements clearly confer strong miR-181 repression. Interestingly, the impact of the repeat-derived MREs was considerably stronger in the translatome compared to the transcriptome (**Figure 5e, f, Figure S6b-h**). Indeed, the transcripts with MMSAT4 MREs showed a significant shift in translational efficiency, indicating that this specific group of transcripts is subject to a distinct mode of miR-181-dependent regulation at the level of translation (**Figure 5g, Figure S6d**).

In conclusion, the comprehensive profiling of miR-181 MREs across transcripts revealed a special class of MREs in the CDS that are associated with satellite repeats and specifically contribute to translational inhibition on top of mRNA destabilization. The identification of a large set of miR-181 targets encoding transcription factors points towards putative new functions of miR-181 and the KZFP family in T-cell development.

## miR-181 seed match complementarity promotes MRE functionality

The binding of miRNA-loaded RISCs to MREs is mediated by different types of seed matches with varying degrees of sequence complementarity (**Figure 6a**). These include the minimal 6mer seed match, involving perfect base pairing between the miRNA seed region (positions 2-7) and the mRNA, a shifted variant 6mer-m8 (positions 3-8) and an extension to 7mer-m8 (positions 2-8). The seed matches 7mer-A1 and 8mer are further expanded by an invariant A nucleotide at position 1 in the mRNA.

**Figure 6.**
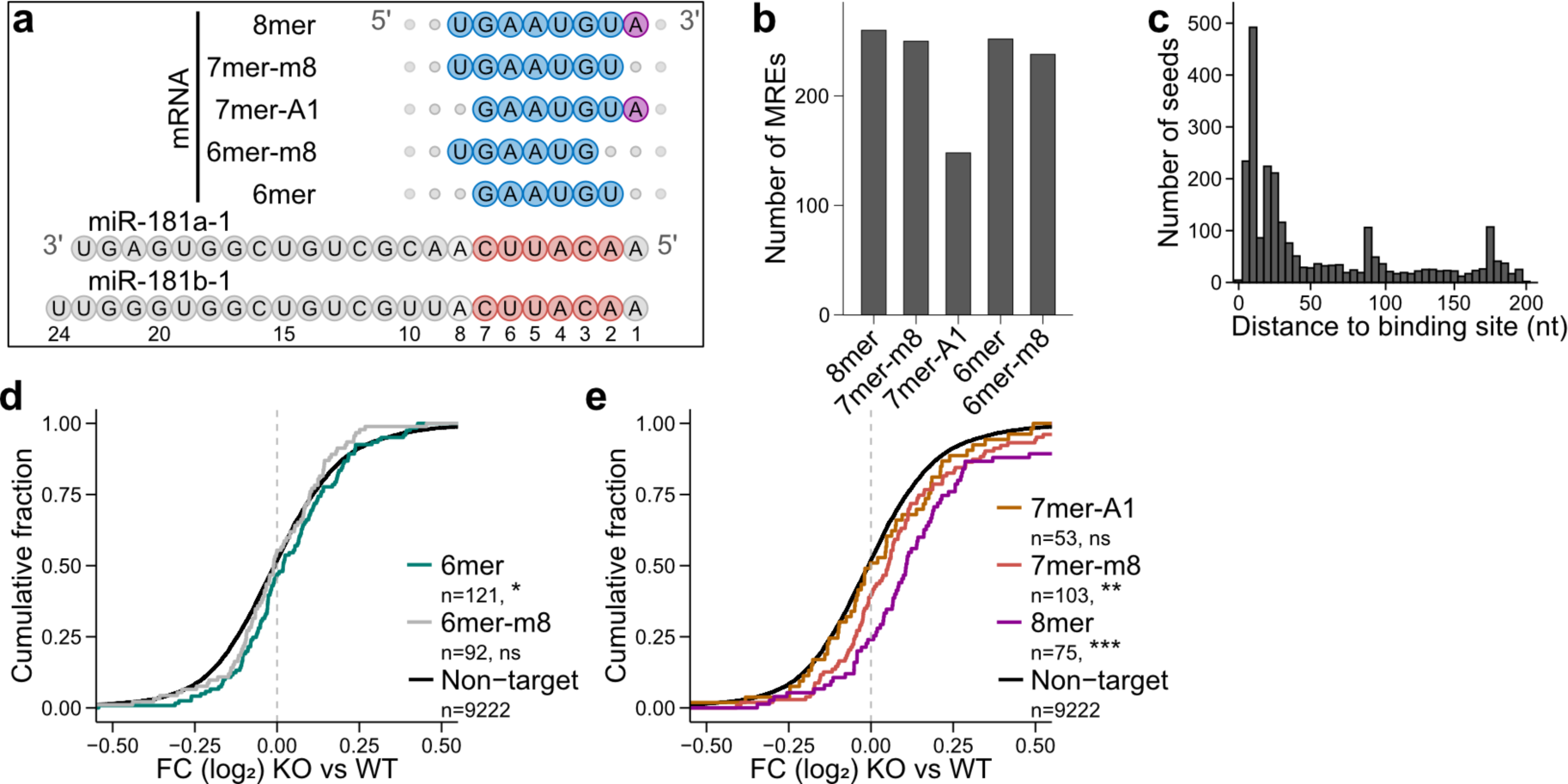
miR-181 seed match complementarity promotes MRE functionality. **(a)** Scheme of different seed match types (top) aligned to the seed sequence in miR-181a and miR-181b sequences (bottom). Seed matches are depicted in blue, the non-complementary A at position 1 is depicted in purple, the seed sequence is depicted in red. **(b)** Number of MREs with different seed match variants. **(c)** Distribution of canonical 6mer seed matches within 200 nt downstream of the binding site. **(d,e)** Fold changes in ribosome occupancy between miR-181 KO and WT conditions are shown as cumulative density of the log_2_ fold changes. Transcripts without MREs are shown in black. (d) Comparison of transcripts with MREs containing 6mer (green) and 6mer-m8 (grey) seed matches. (e) Comparison of 7mer-A1 (orange), 7mer-m8 (red) and 8mer seed matches (purple). Only transcripts with a single MRE were used. Differences between curves were tested for significance using the asymptotic two-sample Kolmogorov-Smirnov test (ns>0.05, *<0.05, **<0.01, ***<0.001).

Using a window up to 200 nt downstream of the MREs, we identified a seed match at 962 MREs (21 %). The different types of seed matches occurred with similar frequency (**Figure 6b**). The majority of seed matches were located within 50 nt downstream of the MREs, underlining that the Ago2 binding sites detected in chimeric eCLIP capture miRNA binding with high resolution (**Figure 6c**). A periodicity of recurring matches further downstream originated from the MREs in satellite repeat elements (see above; **Figure S6a**). We found that all types of seed matches, with the exception of 6mer-m8, were functional in mRNA destabilization. As expected, the repressive capacity correlated with the extent of base-pairing in the seed, with 8mer conferring the strongest repression (**Figure 6d, e, Figure S7a, b**). Consistent with previous reports (9), 7mer-m8 was more active than 7mer-A1, which had comparable effects to 6mer. Taken together, the analysis of MREs confirmed the previously suggested hierarchy of seed matches in miR-181-mediated repression, showing in particular a comparatively weak role of 7mer-A1 matches. However, MREs without seed match still showed a certain repressive capacity, indicating that seed complementarity is not an absolute requirement for regulation.

### A non-canonical miR-181 seed match confers regulation

A *de novo* motif search confirmed the enrichment of seed matches around MREs (**Figure 7a, Figure S8a,b**). Surprisingly, however, the motif indicated the presence of a second variant which contained U instead of A in position 3 of the 6mer seed region to pair with position 5 of miR-181. The alternative seed match GAUUGU (referred to as AU seed match) was present in 507 MREs, thereby reaching almost half the frequency of the canonical GAAUGU (AA seed match, 1,148 MREs; **Figure 7a, b**). Analogous to the canonical AA seed match, the AU seed match occurred in all seed match types (**Figure S7c**) and accumulated within the first 50 nt downstream of the MREs (**Figure 7c, Figure S8c,d**). Intriguingly, across all types, the alternative AU seed match clearly conferred functionality, which was only modestly reduced compared to the canonical variant (**Figure 7d, Figure S7d, e**). In the context of the strongest MREs (8mer), both variants displayed comparable repressive capacity (**Figure 7e, Figure S7f–i**). Dual reporter assays in Scid.adh.2c2 cells ectopically expressing miR-181a showed that triple MREs with a non-canonical seed match displayed a similar repressive capacity as triple MREs containing canonical seed matches (**Figure S5c**). Together, these data indicate that the alternative AU seed match confers a reduced activity, but is fully functional in maximally extended seed matches.

**Figure 7.**
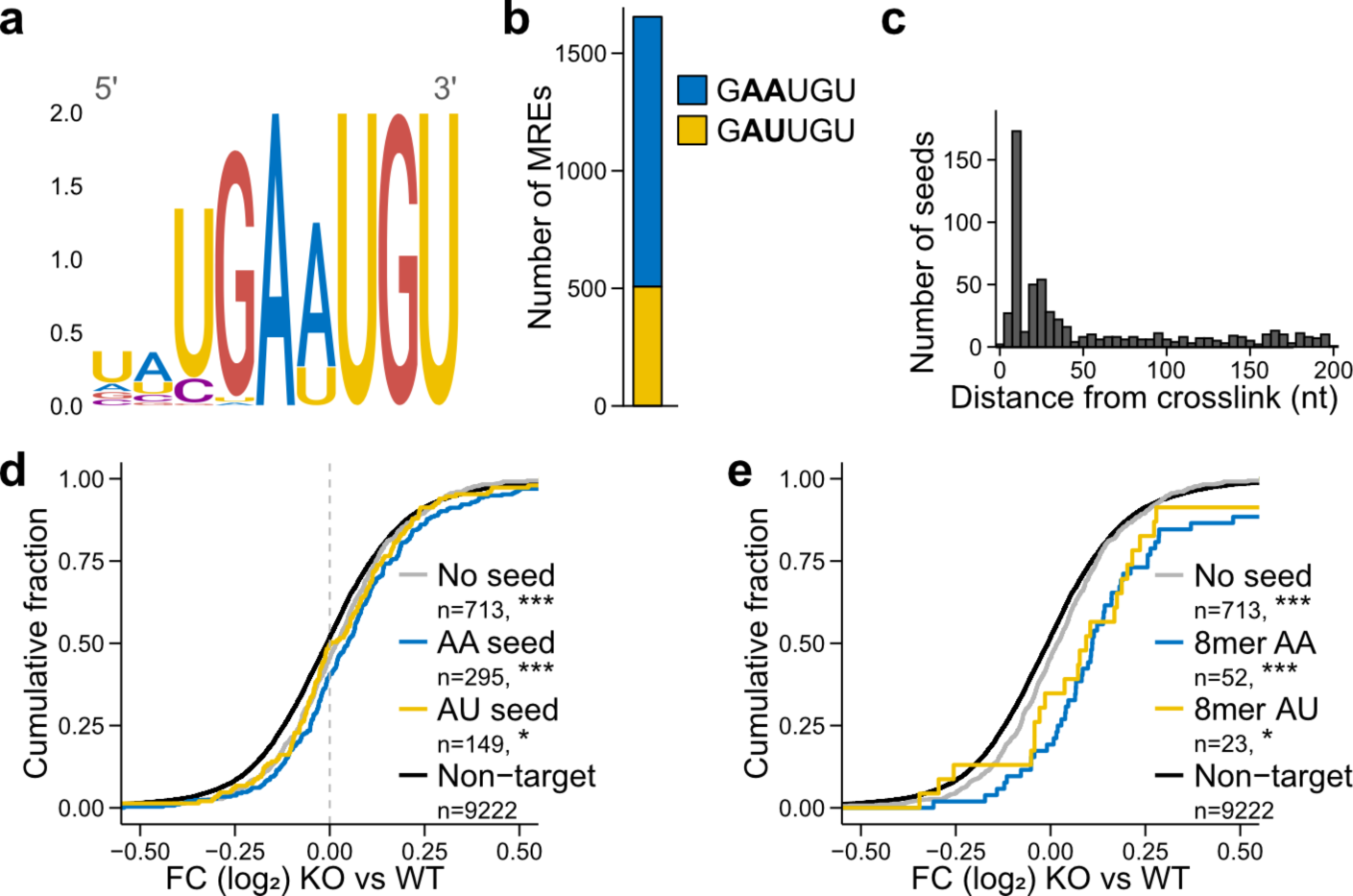
An alternative miR-181 seed match confers regulation. **(a)** Most enriched motif found by *de novo* search in mature transcript sequences within 200 nt downstream of MREs (top) (E-value = 1.6 x 10^-10^). **(b)** Number of MREs containing the canonical 6mer seed match (AA, blue) or the alternative 6mer seed match (AU, yellow) within 200 nt downstream of the MRE. **(c)** Distribution of alternative AU seed matches within 200 nt downstream of the MRE. **(d)** Fold changes in ribosome occupancy between miR-181 KO and WT conditions are shown as the cumulative density of log_2_ fold changes, for transcripts with MREs with the canonical AA seed matches (blue), alternative AU seed matches (yellow), MREs without seed matches (grey) or transcripts without MREs (black). Only transcripts with a single MRE were used. Differences between curves were tested for significance using the asymptotic two-sample Kolmogorov-Smirnov test (ns>0.05, *<0.05, **<0.01, ***<0.001). **(e)** Comparison of canonical 8mer (AA, blue) and alternative 8mer (AU, yellow) to MREs without a seed match. Only transcripts with a single MRE were used. Differences between curves were tested for significance using the asymptotic two-sample Kolmogorov-Smirnov test (ns>0.05, *<0.05, **<0.01, ***<0.001).

In summary, we discovered the unexpected occurrence of a functional A-to-U variant of the seed matches. Together with the observation that seed matches are not absolutely required to mediate repression, these findings support our discovery of a substantially expanded miR-181 targetome when compared to seed-match-centric *in silico* prediction tools.

### Duplex predictions reveal distinct patterns of 3’ base-pairing

Using the identified miR-181 MREs, we investigated the functional role of seed and non-seed base-pairing in miR-181-mediated mRNA repression in thymocytes. To this end, we extracted the sequences from-30 nt to +200 nt around miR-181 MREs and predicted their base-pairing with miR-181a-5p using the algorithm RNAduplex (59) (**Figure 8a, Figure S9**). First, we looked at the sequences that harbored AA seed matches (873 MREs; **Figure S9a, Table S6**). Indeed, many sequences were predicted to form a duplex with miR-181 within this seed match (canonical seed pairing, 408 MREs; **Figure S9b**, top). Nonetheless, a large fraction showed duplex formation outside the seed match (465 MREs; **Figure S9b**, bottom), indicating that neighboring sequence regions compete for binding to miR-181. Consistently, MREs with competing duplex formation outside of the seed match were associated with a lower repressive efficacy (**Figure S9c**). For the MRE sequences lacking a genuine seed match, RNAduplex predicted two scenarios in which the seed region of miR-181 was either partially paired (2,229 MREs) or completely devoid of base-pairing (no seed pairing, 665 MREs, **Figure 8a, Figure S9d**). Using RF data as a functional readout, we found that MREs with either scenario still had repressive capacity, which was reduced compared to MREs with canonical seed pairing but comparable between MREs with partial seeds or no seed pairing (**Figure 8b**).

**Figure 8.**
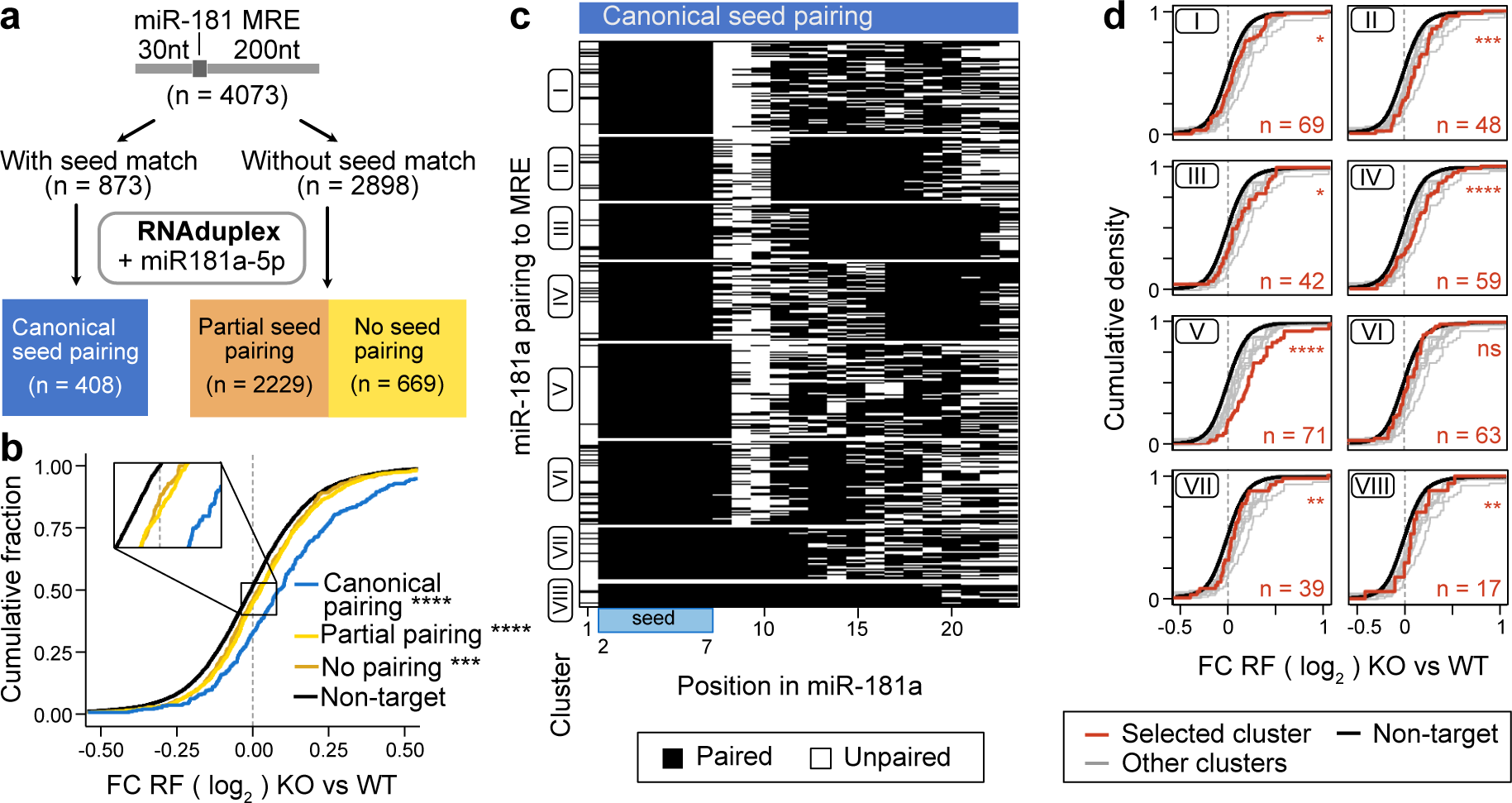
Duplex predictions reveal distinct patterns of 3’ base-pairing. **(a)** Scheme of the base-pairing analysis. Duplex formation with miR-181a was predicted with RNAduplex using the mature transcript sequences in a window from 30 nt before until 200 nt after the miR-181 MREs. Target regions containing a seed match (n = 873) were analyzed separately from those without a seed match (n = 2898). Targets without seed match were further split between duplex structures with some (n = 2229) or no (n = 669) pairing within the first 8 nt of the miR-181a sequence. **(b)** Fold changes in ribosome occupancy between miR-181 KO and WT conditions are shown as cumulative density of log_2_ fold changes. Comparison of transcripts with canonical seed pairing (blue), partial seed pairing (yellow) or no seed pairing (orange). Transcripts without MREs are shown in black. **(c)** Heatmap of miR-181a base-pairing in duplexes with MRE sequences with canonical seed pairing. Structures were clustered by *k*-means clustering (*k* = 8). White – unpaired, black – paired. Blue bar at the bottom indicates the seed region. **(d)** Fold changes in ribosome occupancy between miR-181 KO and WT conditions are shown as cumulative density of log_2_ fold changes. Comparison of clusters for canonical duplex formation. The respective cluster is colored in red, and the number of transcripts given at the bottom right. All other clusters are colored in grey. Transcripts without MREs are shown in black. Differences between curves were tested for significance using the asymptotic two-sample Kolmogorov-Smirnov test (ns>0.05, *<0.05, **<0.01, ***<0.001, ****<0.0001).

Next, we analyzed the occurrence of additional 3’ binding in MREs with canonical seed pairing. Clustering revealed various patterns of additional base-pairing (**Figure 8c, Figure S9e,h**). Consistent with structural constraints within the RISC complex, six out of eight clusters were devoid of base-pairing around position 10 of the miRNA (69). However, two clusters showed extensive consecutive base-pairing up to position 12 (cluster VII) and 18 (cluster VIII), respectively. Furthermore, 3’ base-pairing was identified around positions 11-15 and/or 17-19 (clusters V, VI), 11-17 (cluster II), 13-21 (cluster III) or 17-21 (cluster IV). Analyses of the repressive capacity showed a strong component of seed binding length (cf. cluster V vs. cluster I), with highest repressive capacity in cluster V, which almost exclusively contain 7mer-m8 or 8mer seed matches (**Figure 8d**). 3’ supplemental pairing increased the repressive capacity of short seed matches (clusters II, III, IV vs. cluster I), although this effect might be partially due to some extended seed matches in these clusters. Surprisingly, 3’ supplemental pairing limited the efficacy of repression of long-seed MREs (cluster VI vs. cluster V), possibly due to the introduction of structural constraints. However, long consecutive stretches of base-pairing (cluster VII, VIII) did not fully abolish repression.

### Duplex prediction reveals functional non-canonical base pairing in miR-181 MREs

Similar to the MREs with canonical base-pairing, MREs with partial seed matches showed various patterns of additional 3’ base-pairing at positions 17-22 (cluster I), 12-17 (cluster IV) or 12-21 (cluster VI) (**Figure 9a, Figure S9f,i**). Clusters I and VI had moderate repressive capacity, similar to cluster II, which lacked defined contiguous 3’ base-pairing (**Figure 9b**). We also detected two clusters corresponding to so-called centered sites, of which only cluster III with a shorter stretch of contiguous base-pairing had repressive capacity (16).

**Figure 9.**
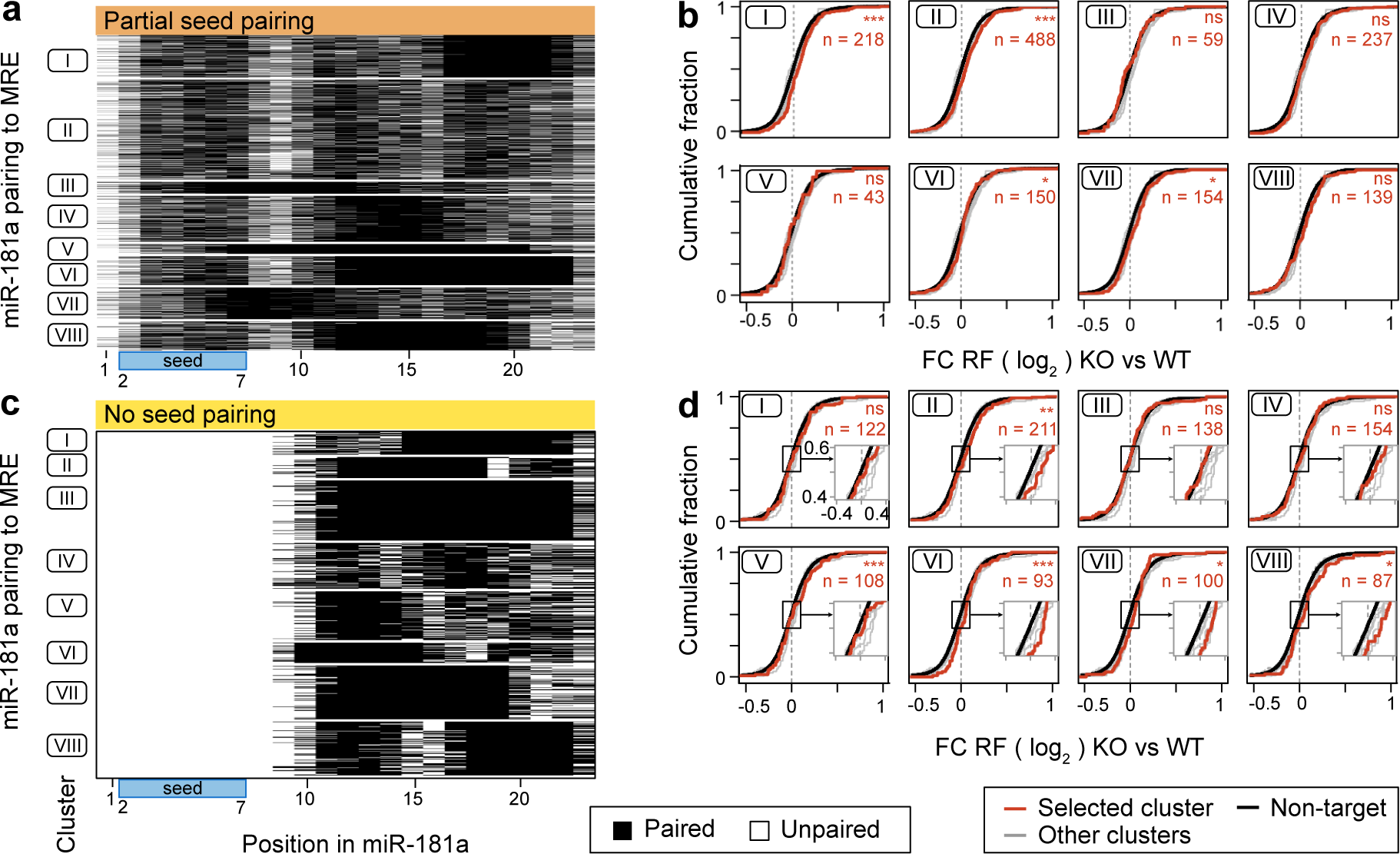
Duplex prediction reveals functional non-canonical base pairing in miR-181 MREs. **(a,c)** Heatmap of miR-181a base-pairing in duplexes with MRE sequences with partial seed pairing (a) and no seed pairing (c). Structures were clustered by *k*-means clustering (*k* = 8). White – unpaired, black – paired. Blue bar at the bottom indicates the seed region. **(b,d)** Fold changes in ribosome occupancy between miR-181 KO and WT conditions are shown as cumulative density of log2 fold changes. Comparison of clusters for partial seed pairing (b), or no seed pairing (d). The respective cluster is colored in red, and the number of transcripts given at the bottom right. All other clusters are colored in grey. Transcripts without MREs are shown in black. Differences between curves were tested for significance using the asymptotic two-sample Kolmogorov-Smirnov test (ns>0.05, *<0.05, **<0.01, ***<0.001).

Finally, we investigated MREs without seed pairing, i.e., completely lacking complementarity around miRNA positions 2-7 in the predicted duplex structures. MREs without seed pairing showed extensive 3’ binding with distinct patterns containing either 1 (clusters I, III, V, and VII) or 2 (clusters II, IV, VI, and VIII) stretches of contiguously paired bases (**Figure 9c, Figure S9g,j**). The base-pairing characteristics of these MREs did not strongly correlate with repressive capacity (**Figure 9d**). However, MREs with short stretches of contiguous base-pairing (cluster IV) or, conversely, near-complete base-pairing throughout the 3’ region (cluster III), were not conducive to repression.

Taken together, these analyses indicated that seed complementarity is a strong, but non-exclusive predictor of functional MREs of miR-181 in primary thymocytes. Of note, 3’ supplemental pairing increased the repressive capacity of MREs with short seed matches. Seed match-deficient MREs were mostly active when harboring two stretches of contiguous base-pairing. Thus, various distinct 3’ base-pairing patterns expand the pool of potentially relevant targets of miR-181 in thymocytes.

## Discussion

miR-181a/b-1 constitutes one of the most prominently expressed miRNA clusters in the murine thymus (25,24). Functionally, miR-181 is critical for the development of unconventional T cells and has also been implicated in the selection of conventional T cells (29,70,27,31,32,30). Like the majority of miRNAs, miR-181 has a large number of predicted targets, and several of them have been implicated in the molecular mechanisms by which miR-181 governs T-cell selection.

Here, we took advantage of the high levels of miR-181 in thymocytes and the availability of genetic deletion models which enabled us to identify and characterize functional miR-181 targets without ectopic expression in primary cells. To achieve this, we combined deep profiling of miR-181 MREs in primary murine thymocytes using chimeric eCLIP with their functional characterization using differential translatome and transcriptome analysis of miR-181a/b-1 KO and WT cells. We identified MREs in numerous previously validated targets, including *Dusp6*, *Ptpn22*, *Bcl2*, *Cd69*, and *S1pr1* (24,31,32). We also found a large number of novel targets. Among these are RNA-binding proteins of the Zfp36 family, Zfp36l1 and Zfp36l2, which have been implicated in controlling early T-cell development through regulation of the cell cycle and Notch signaling (63,64). Interestingly, GO term enrichment revealed MREs in transcripts associated with various processes of global gene regulation, including chromatin modification, RNA processing and miRNA-mediated post-transcriptional gene regulation itself. A similar miRNA-dependent negative feedback on RISC levels has been recently proposed for the miR-182/miR-183 pair in embryonic stem cells (71). Targeting of global gene regulatory modules, including modifiers of chromatin accessibility, may also provide a mechanistic explanation for the observation that complete deletion of all three miR-181 clusters is incompatible with life (30).

Analysis of miR-181 MREs revealed a distinct periodicity of seed matches in a number of mRNAs, which originated from satellite repeat elements of the MMSAT4 and MurSatRep1 families. Whereas little is known about the latter, the MMSAT4 elements are associated with genes encoding for C2H2 zinc finger families, in particular KRAB-containing poly-zinc finger proteins (KZFPs), which form the largest family of transcription factors in mammalian genomes (68). Consistent with prior reports employing *in silico* prediction and overexpression in cell lines, we found that in contrast to the majority of MREs, MREs associated with MMSAT4 are primarily located in CDS and mediate strong repression (72,73). It is likely that the conservation of miR-181 MREs in the CDS of C2H2 zinc finger family members is directly related to the fact that one cysteine-encoding base triplet is complementary to a part of the miR-181 seed (72). It has been proposed that many KZFPs act as transcriptional repressors and are likely to target transposable elements. Given that the origin of satellite repeats lies in such transposable elements, it is interesting to speculate that miR-181 modulates this feedback regulation (68). Our study biochemically validates this unusual group of miR-181 targets in primary cells and uncovers that this pool of targets is selectively regulated by translational inhibition on top of RNA destabilization. This suggests that translational inhibition in this case may be a feature of miRNA-mediated gene regulation, when binding of multiple RISCs induces ribosome stalling on mRNA, rather than competition of RISC components with initiation factors for cap binding (3–5). Notably, base-pairing analysis revealed no common composition of this subgroup of MREs, as has been proposed by others (5). Rather, we observed a similar distribution of base-pairing patterns in repeat MREs as identified in the complete pool of MREs. Our finding also provides an interesting evolutionary perspective, in which C2H2 transcription factors, which have been proposed to transcriptionally suppress their retroviral origin, have co-evolved with their own means of negative control, allowing for selective downregulation of KZFPs when levels of miR-181 are high. It will be interesting to further characterize the miR-181–KZFP targeting network, as the role of KZFP transcription factors has not been explored in the context of T-cell development. Given the repetitive nature of MREs in these transcripts, it is reasonable to speculate that they could act as “miRNA sponges” as well, providing an additional rheostat of miR-181 abundance (74).

By chimeric eCLIP (60), we faithfully identified more than 6,000 miR-181 MREs in more than 2,000 mRNAs in primary thymocytes for an individual miRNA. This corresponds to a five-fold increase over our initial Ago2-IP without miRNA enrichment and represents a substantial expansion of the MRE pool for an individual miRNA compared to previous studies (75,12,13). Sensitive mapping of miR-181 MREs in combination with differential translatome and transcriptome data allowed us to characterize the nucleotide-level composition of functionally active MREs in primary thymocytes directly *ex vivo* at high-resolution. Importantly, the number of mRNAs containing a single MREs was large enough to probe individual features of MREs for efficacy. Our data support canonical models of seed-based miRNA-mediated repression, including a biochemically established hierarchy of site efficacy of 8mer > 7mer-m8 > 7mer-A1 > 6mer (9–11). However, only about 30% of MREs contained a seed match of any kind. Although MREs with incomplete or no seed match were less effective overall, we found a substantial contribution to miR-181-mediated repression from those sites as well.

Given the large number of mR-181 MRE available for analysis at nucleotide level, we were also able to investigate the contribution of 3’-end base pairing. Distinct 3’ base pairing patterns improved the efficacy to minimal 6mer seed match MREs, but to a lower extent than seed match extension, supporting the dominant role of the seed region in miRNA-mediated targeting. In turn, contiguous base pairing through positions 9-11 of the miRNA limited seed efficacy, presumably due to structural constraints imposed by the RISC complex (69,76). 3’-base pairing was extensive in the absence or partial absence of seed matches with distinct base pairing patterns correlated with varying efficacies of repression. Interestingly, the length of contiguous pairing was not associated with repressive capacity. Surprisingly, we identified a frequent A-to-U mismatch in seed matches for miR-181, which was also associated with substantial repressive capacity. Taken together, nucleotide-level analysis of miR-181 MREs revealed extensive base pairing beyond the seed, similar to miRNA–mRNA binding patterns described before (13,12). However, larger numbers of MREs identified for an individual miRNA pair allowed for more refined stratification and functional testing.

Information on the functional targetome of individual miRNAs based on biochemical characterization using CLIP-like approaches or direct genetic interference with individual miRNA–MRE interactions in the immune system remains scarce. Deletion of individual MREs has shown that *in silico* target prediction, even in combination with functional validation in overexpression systems, may reveal highly relevant, partially functional or non-functional MREs (77–79). Such approaches currently lack the potential to uncover co-targeting networks, a feature of miRNAs associated with their capacity to target many mRNA with mild repressive effects on a per mRNA basis (80). Differential CLIP analysis of miR-155 in WT vs KO mice revealed important contextual cell-type information on the miR-155 targetome in immune cells (81). Our study extends this type of analysis by providing substantially increased depth and the possibility of nucleotide-level characterization of functionally relevant MREs. Thus, we were able to uncover the abundance and features of non-canonical MREs, and the alternative seed match in particular, considerably expanding the currently predicted targetome of miR-181. Hence, our study provides a valuable resource for the characterization of miR-181 function in the immune system as well as novel information on the patterns of biochemically identified and functionally active MREs *in vivo* (20).

### Limitations of the study

Throughout this study, total murine thymocytes were utilized, as ribosome footprinting relies on immediate *ex vivo* fixation of ribosomes on mRNAs, and all other datasets were chosen to maintain consistency with this condition. As a consequence, the miR-181 effects may be partially confounded by the varying abundance of thymocyte populations, including iNKT cells, MAIT cells and Treg cells, which are reduced in the absence of miR-181 (70,29,27). However, miR-181 is predominantly expressed in double-positive thymocytes, which are unaltered in number in KO thymi. Identification of MREs by chimeric eCLIP is, therefore, almost exclusively restricted to these cells and mRNAs not found in double-positive thymocytes, are classified as non-targets throughout. Additionally, CLIP protocols inherently favor detecting protein binding sites on highly abundant transcripts, while lowly expressed transcripts are typically disadvantaged (82). Moreover, overlapping transcript isoforms and conflicting annotations mean that not all detected binding sites can be unambiguously assigned to target transcripts. Hence, any limitations in this regard likely lead to an underestimation of the observed effects.

## Data availability

All sequencing data is available in the Gene Expression Omnibus (GEO) under the SuperSeries accession number GSE242306. The collection includes the RNA-seq and Ribosome footprinting data from WT and miR-181a/b-1-deficient thymocytes (GSE242304) as well as the miR-eCLIP data (GSE241905). Code generated in the context of this study can be found at https://zenodo.org/doi/10.5281/zenodo.10863700. The `racoon_clip` pipeline is available at https://github.com/ZarnackGroup/racoon_clip. A detailed description of the usage and functions of the miR-eCLIP in `racoon_clip` and is available at: https://racoon-clip.readthedocs.io/en/latest/tutorial_mir.html. miR-eCLIP and RF data can be visualized at https://genome.ucsc.edu/s/melinak/miR181%2DeCLIP_Verheyden%26Klostermann_2024 and https://genome.ucsc.edu/s/melinak/miR181_RibosomeFootprint_Verheyden%26Klosterman <u>n_2024</u>, respectively.

## Acknowledgements

We are grateful to all members of the Krueger and Zarnack labs for helpful discussions and to Julia E. Weigand, Marburg, for the gift the pDF-FRT dual reporter vector. We thank Julia Häusler for mouse colony management.

## Author contributions

N.A.V., A.S., S.A., C.L., H.M.S. performed experiments and analyzed data. M.K., M.B. developed bioinformatic analysis pipelines and analyzed data. C.M., T.S., K.Z., A.K. designed studies and interpreted data. N.A.V., M.K., K.Z., A.K., wrote the manuscript and made the figures.

