## Supplementary Information for "A high-resolution map of functional miR-181 response elements in the thymus reveals the role of coding sequence targeting and an alternative seed match"

### Supplementary Figures

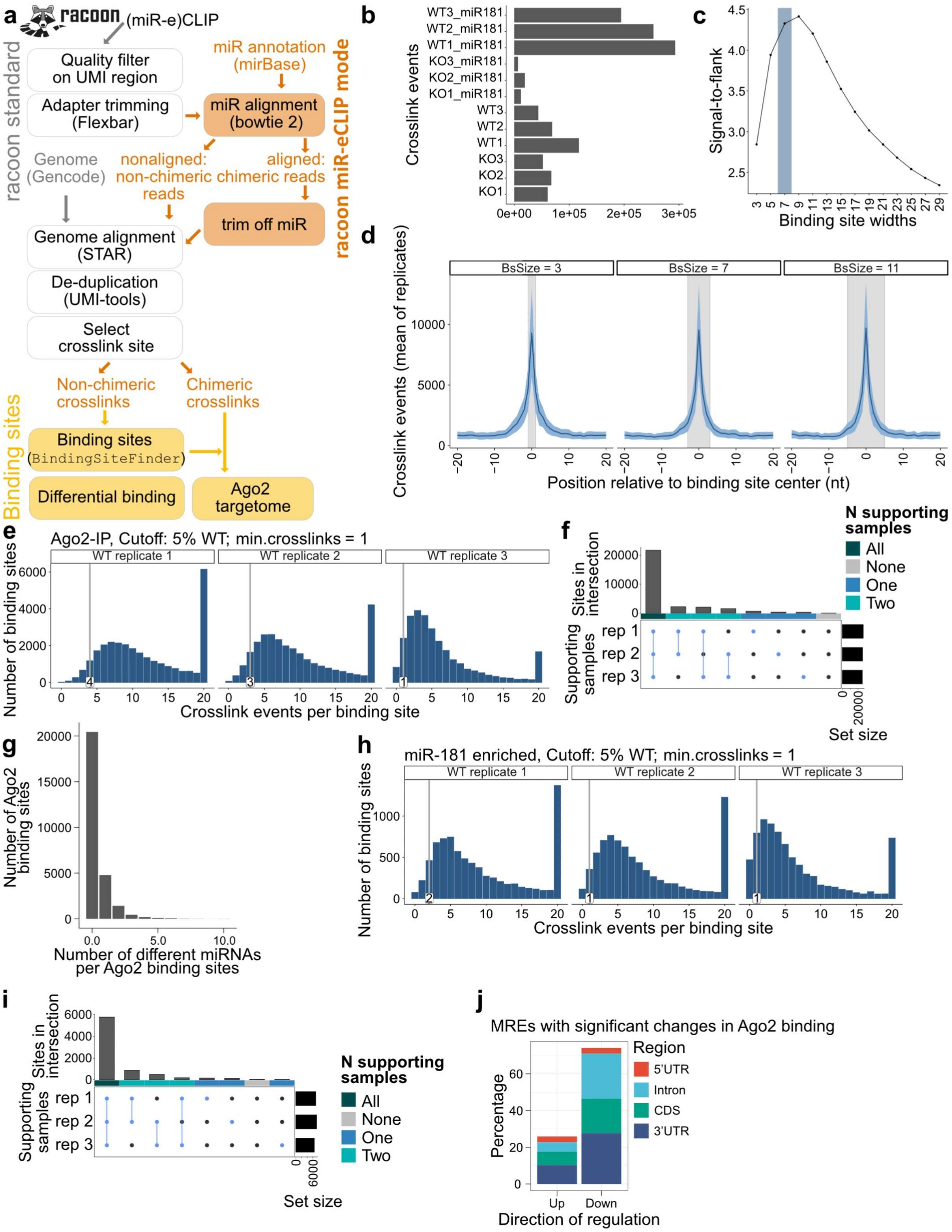


**Figure S1. Identification of reproducible Ago2 binding sites from chimeric eCLIP data. (a)** Processing of miR-eCLIP data with `racoon` and `BindingSiteFinder`. Non-chimeric reads are processed like standard eCLIP data (grey). Chimeric miR-eCLIP reads are aligned to the miRNA annotation. The chimeric part of the read is then trimmed off and the leftover read is treated like the non-chimeric reads (orange). After `racoon` processing,

`BindingSiteFinder` is used to define binding sites on the different subsets. These binding sites are then also further used for differential binding and together with the chimeric reads to define the AGO targetome. **(b)** Number of crosslink events per miR-eCLIP sample. miR-eCLIP was performed on the four different conditions with three replicates each: crosslinks from Ago2-IP (WT), crosslinks from Ago2-IP (KO), and crosslinks from miR-181a/b enrichment after AGO-IP without and with miR-181 KO. **(c, d)** Determination of optimal binding site width using (c) signal-to-flank ratio for binding sites from 3 to 29 nt width, and (d) distribution of crosslink events in the proximity of binding sites for binding sites of 3, 7 or 11 nt widths. **(e, f)** Reproducibility filtering for Ago2 binding sites. (e) A sample-wise cutoff was defined at the lowest 5% quantile of crosslinks per binding sites. (f) Upset plot shows the number of binding sites supported by each sample after applying the 5% cutoff. Number of supporting samples is displayed by colored bars above (dark green – all, cyan – two, blue – one, grey – none). Only binding sites supported by all three replicates were deemed reproducible and kept for further analysis. **(g)** Distribution of chimeric reads per Ago2 binding site. **(h, i)** Reproducibility filtering for miR-181 MREs. (h) A sample-wise cutoff was defined at the lowest 5% quantile of crosslinks per binding sites. (i) Upset plot shows the number of binding sites supported by each sample after applying the 5% cutoff. Number of supporting samples is displayed by colored bars above (dark green – all, cyan – two, blue – one, grey – none). Only binding sites supported by all three replicates were deemed reproducible and kept for further analysis. **(j)** The number of significantly up- and downregulated binding sites in miR-181 KO cells are shown colored by the gene regions assigned to the binding sites (red - 5'UTR, light blue – intron, green – CDS, dark blue – 3'UTR).

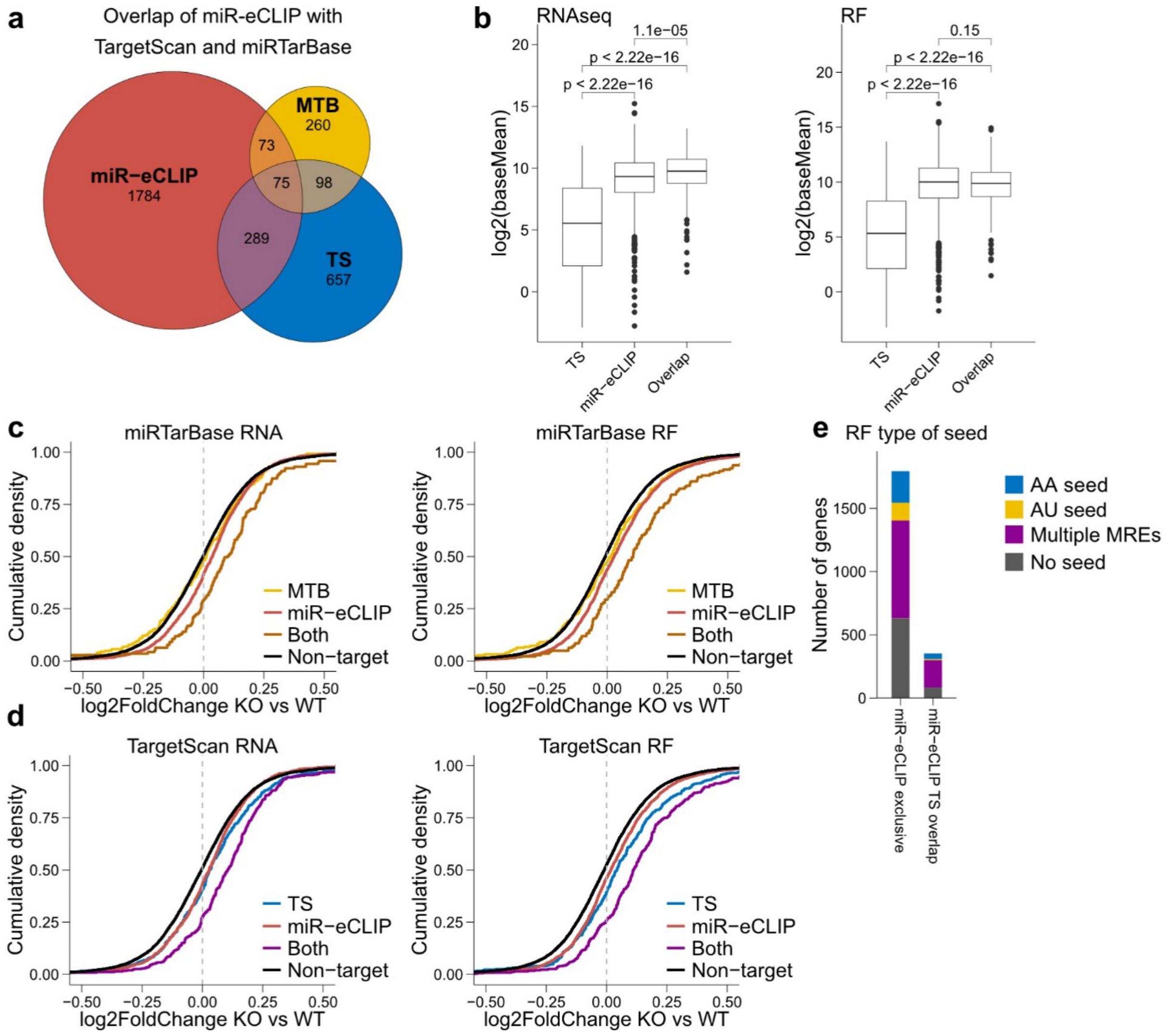


**Figure S2. Comparison of miR-181 target sets from miRTarBase, TargetScan and this study. (a)** Overlap of miR-181 target genes identified in miR-eCLIP (this study), miRTarBase and TargetScan. **(b)** Mean normalized reads (log2) of RNAs in RNAseq (left) and RF (right). TS, TargetScan predictions alone; miR-eCLIP, this study; overlap, overlap as in (a). **(c)** Fold changes in transcript levels (left) or ribosome occupancy (right) between miR-181 KO and WT conditions are shown as the cumulative density of the $\log_2$ values, for transcripts listed in miRTarbase alone (MTB), this study alone (miR-eCLIP), both or no miR-181 MREs. **(d)** Fold changes in transcript levels (left) or ribosome occupancy (right) between miR-181 KO and WT conditions are shown as the cumulative density of the $\log_2$ values, for transcripts listed in TragetScan alone (TS), this study alone (miR-eCLIP), both or no miR-181 MREs. **(e)** Quantification of genes according to seed match type and number of MREs. miR-eCLIP, this study; miR-eCLIP-TS overlap, transcripts identified in this study and listed in TargetScan.

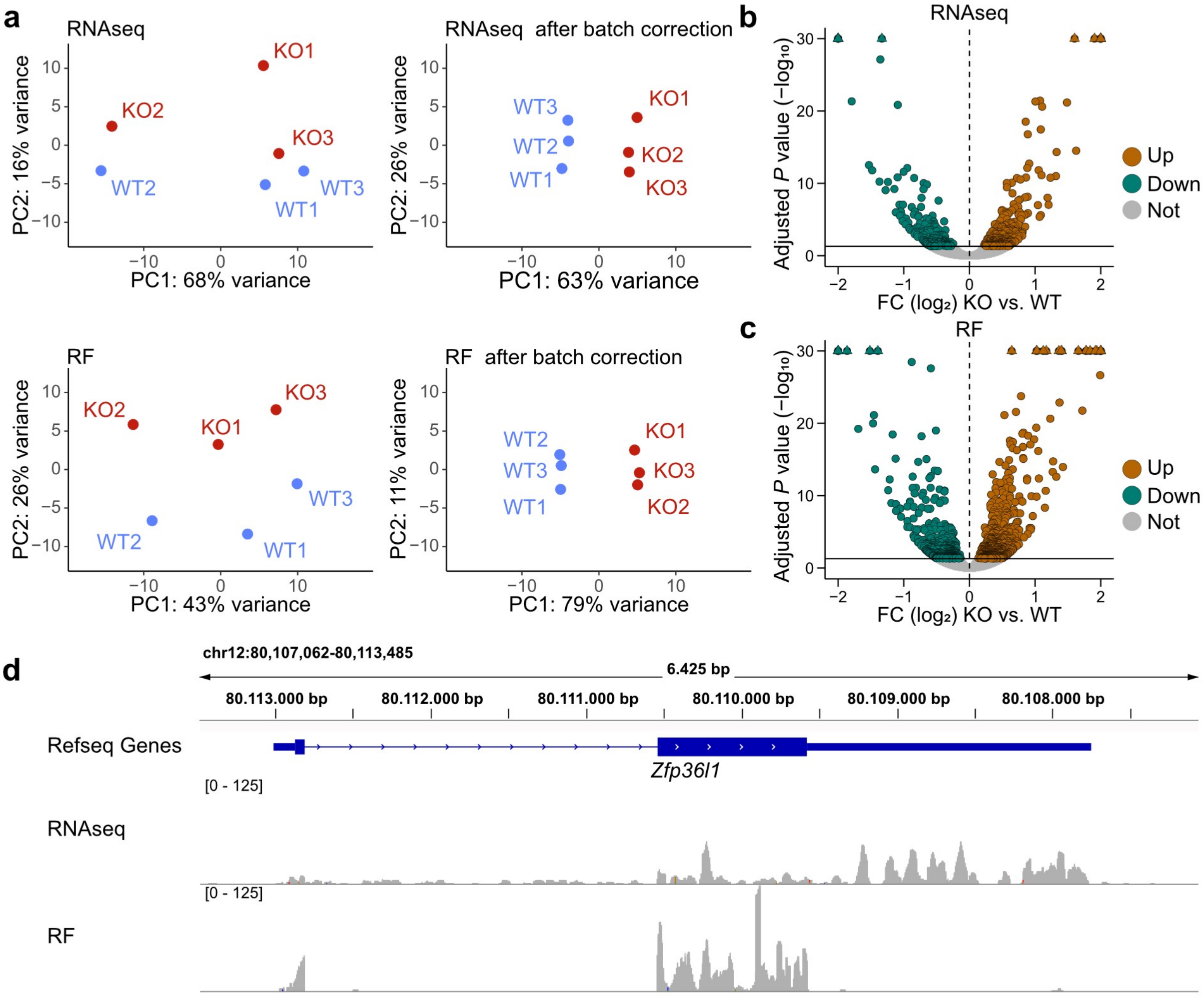


**Figure S3. RNAseq and ribosome footprinting (RF) reveal genes in gene expression upon miR-181 KO. (a)** Principal component analysis of biological replicates of RNAseq (top) and RF (bottom) data before (left) and after (right) batch correction with CombatSeq. **(b,c)** Volcano plots of RNAseq and RF data shown in Figure 3b,c plotted as $\log_2$ fold change (FC) of KO over WT vs. adjusted *P*-value ($-\log_{10}$). **(d)** Genome browser view of RNAseq (top) and RF (bottom) data in *Zfp36l1*.

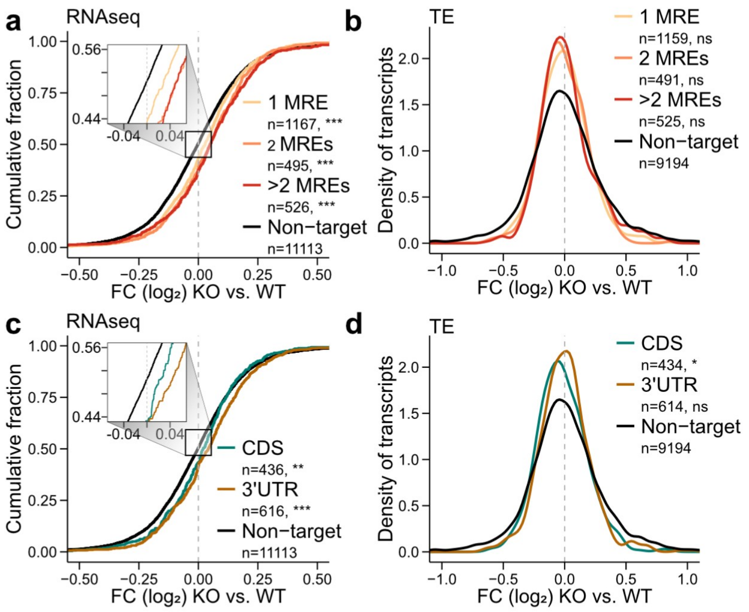


**Figure S4. Effects of MRE number and positioning on transcript abundance (RNAseq) and translational efficiency (TE). (a)** Fold changes in transcript levels between miR-181 KO and WT conditions are shown as the cumulative density of the $\log_2$ values, for transcripts with one (yellow), two (orange), more than two (red) or no miR-181 MREs (black). Box on the left depicts zoom-in. Numbers of transcripts in each set are given. Asterisks indicate the significance of difference to genes without MREs (ns>0.05, *<0.05, **<0.01, ***<0.001, asymptotic two-sample Kolmogorov-Smirnov test). **(b)** Changes in translational efficiency (TE) for transcripts with MREs vs. non-targets shown as $\log_2$ fold-changes of KO over WT vs. density, for transcripts with one (yellow), two (orange), more than two (red) or no miR-181 MREs (black). Numbers of transcripts in each set are given (ns>0.05, *<0.05, **<0.01, ***<0.001, Welch two sample t-test). **(c)** Fold changes in transcript levels between miR-181 KO and WT conditions are shown as the cumulative density of the $\log_2$ values, for transcripts with MREs in the CDS (yellow), 3' UTR (blue), or without MREs (black). Numbers of transcripts in each set are given. Box depicts zoom-in. Only transcripts with a single MRE were used. Asterisks indicate the significance of difference to genes without MREs (ns>0.05, *<0.05, **<0.01, ***<0.001, asymptotic two-sample Kolmogorov-Smirnov test). **(d)** Changes in translational efficiency (TE) for transcripts with MREs vs. non-targets shown as $\log_2$ fold-changes of KO over WT vs. density, for transcripts with MREs in the CDS (yellow), 3' UTR (blue), or without MREs (black). Numbers of transcripts in each set are given (ns>0.05, *<0.05, **<0.01, ***<0.001, Welch two sample t-test).

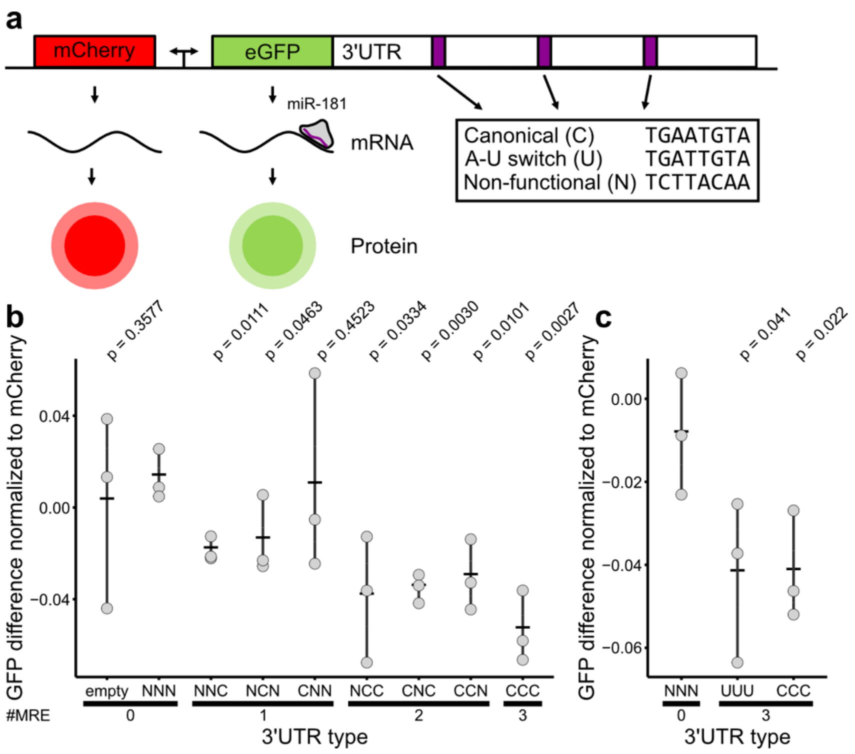


**Figure S5. Reporter assay to test impact of MRE number and alternative seed match. (a)** Scheme of reporter construct design, with mCherry as expression control and 3' UTRs containing 3 identical or varying miR-181 MRE types fused to the coding sequence of eGFP. The ratio of eGFP/mCherry fluorescence serves as readout for miR-181-medated repression. **(b)** Analysis of the effect of MRE number and position on repressive capacity. Three letter code indicates the order of MRE types in the synthetic eGFP 3' UTR (C, canonical; N, non-functional). Empty indicates the eGFP/mCherry ratio in the absence of the synthetic 3' UTR. **(c)** Analysis of the effect of non-canonical (UUU) versus canonical (CCC) MREs on repressive capacity. NNN: non-functional MREs. Data are from 3 independent experiments. *P* values from one-tailed *t* test.

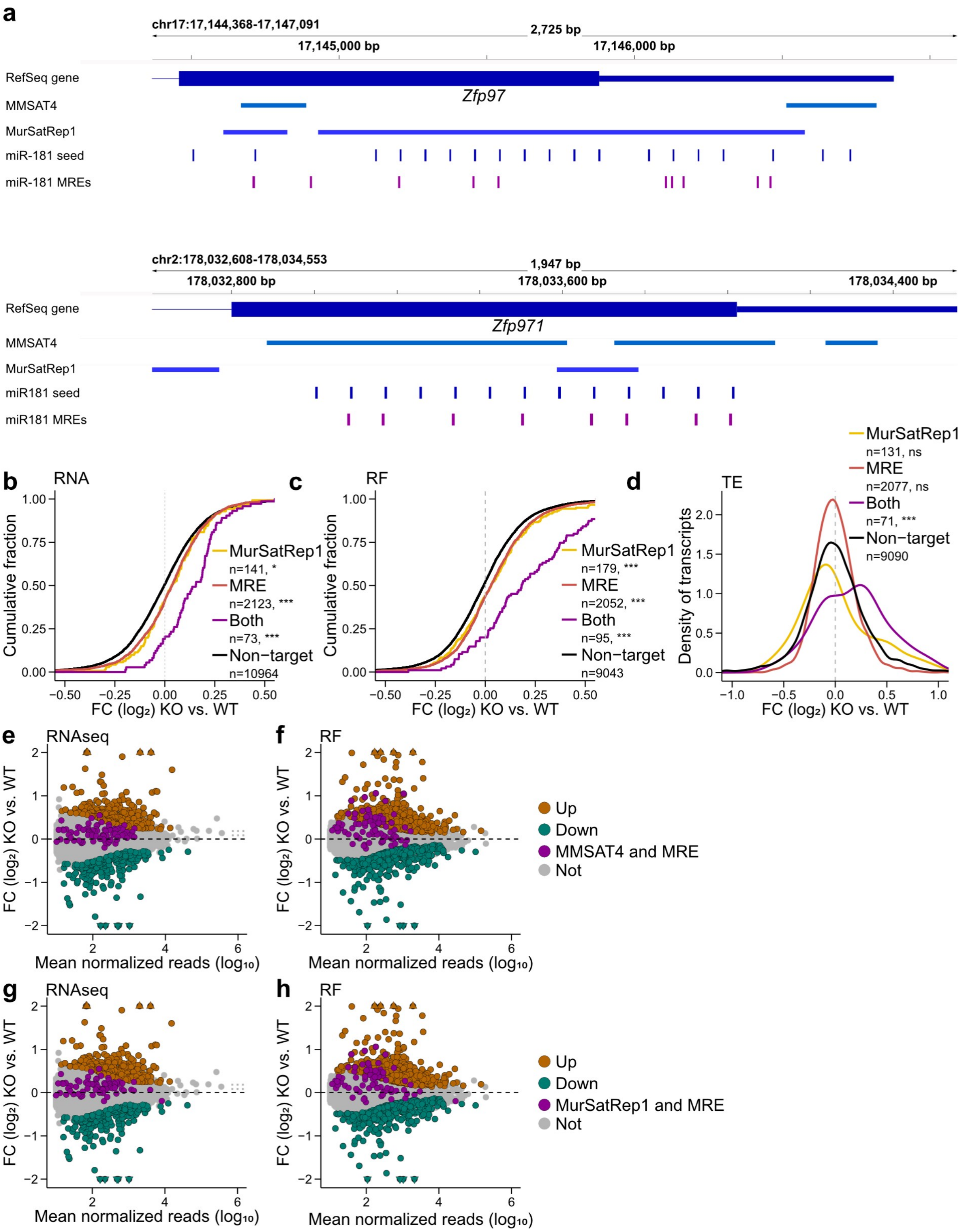


**Figure S6. miR-181 MREs accumulate in satellite repeats. (a)** Distribution of MMSAT4 and MurSatRep1 repeats, miR-181 seed matches, and miR-181 MREs in transcripts of the two representative zinc finger protein (Zfp)-encoding genes *Zfp97* and *Zfp971*. **(b–d)** Fold changes in transcript levels (b), ribosome occupancy (c) are shown as cumulative density of the $log_2$ values for transcripts containing MurSatRep1 repeats (yellow), transcripts containing MREs (red), transcripts containing both MREs and MurSatRep1 repeats (purple) and not containing

MREs or MurSatRep1 repeats (black); Asterisks indicate the significance of difference to transcripts without MREs (ns>0.05, *<0.05, **<0.01, ***<0.001, asymptotic two-sample Kolmogorov-Smirnov test). (d) Translational efficiency is shown as density (ns>0.05, *<0.05, **<0.01, ***<0.001, Welch two sample *t*-test). **(e–h)** Transcripts with significant differences (adjusted *P* value < 0.05, Benjamini-Hochberg correction) in RNA abundance (RNAseq) (e,g) or ribosome occupancy (RF) (f,h) are plotted as log2 fold change (FC) of KO over WT vs. mean normalized read count as shown in Figure 3b,c. transcripts containing both MREs and MMSAT4 repeats (e,f) or MREs and MurSatRep1 repeats (g,h) are highlighted in purple.

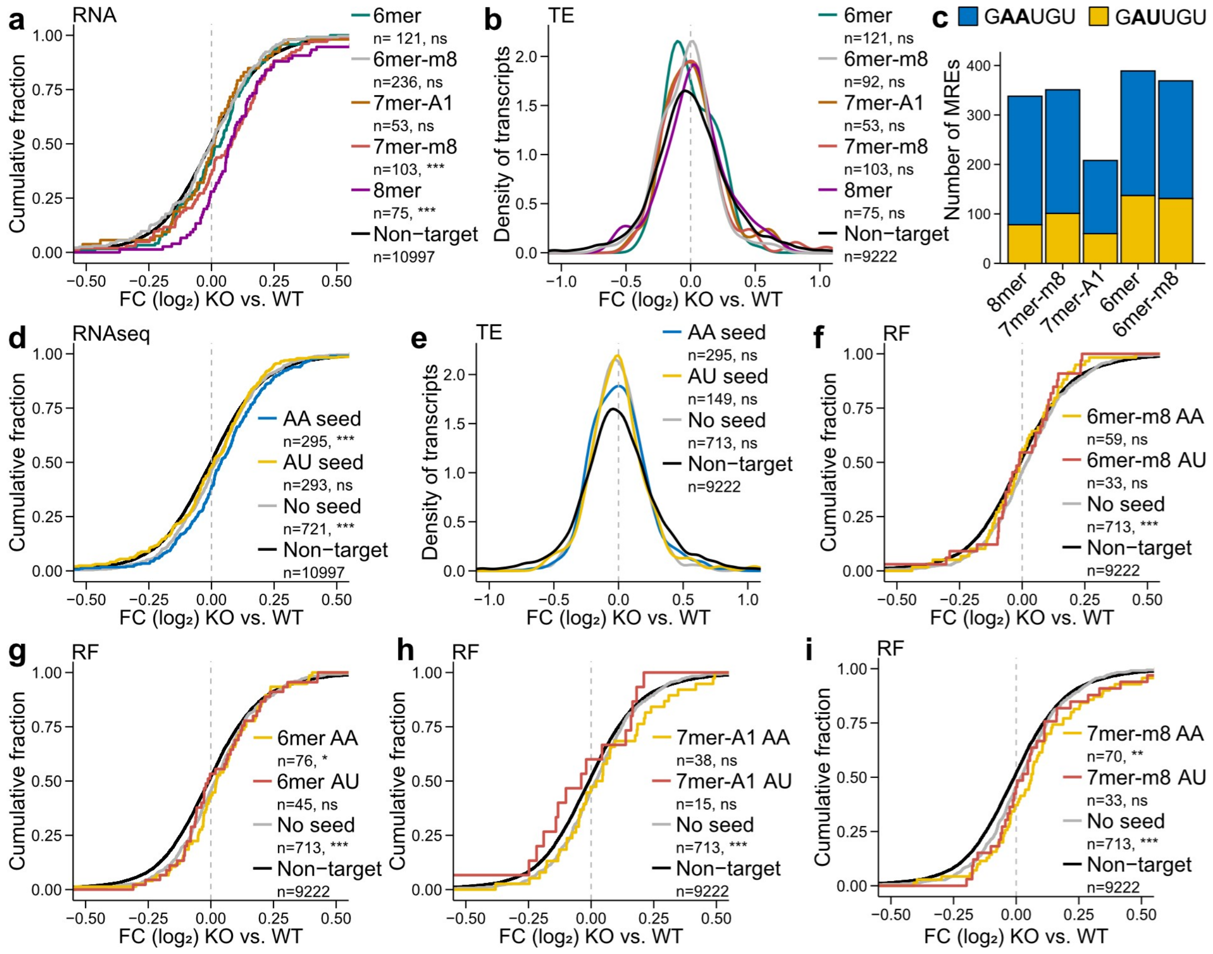

**Figure S7. Effects of different seed matches on ribosome footprinting (RF), transcript abundance (RNAseq) and translational efficiency (TE). (a)** Fold changes in transcript levels between miR-181 KO and WT conditions are shown as the cumulative density of the $log_2$ values. Comparison of transcripts with MREs containing 6mer (green), 6mer-m8 (grey), 7mer-A1 (brown), 7mer-m8 (red) and 8mer seed matches (purple). Numbers of transcripts in each set are given. Asterisks indicate significance of difference to genes without MREs (ns>0.05, *<0.05, **<0.01, ***<0.001, asymptotic two-sample Kolmogorov-Smirnov test). **(b)** Changes in TE for transcripts with MREs vs. non-targets shown as $log_2$ fold-changes of KO over WT vs. density. Comparison of transcripts with different MREs as in (a). Numbers of transcripts in each set are given (ns>0.05, *<0.05, **<0.01, ***<0.001, Welch two sample *t*-test). **(c)** Quantification of MREs according to seed match type. **(d)** Fold changes in transcript levels between miR-181 KO and WT conditions are shown as the cumulative density of the $log_2$ values. Comparison of transcripts with MREs with the canonical AA seed match sequence (blue), the alternative AU seed match (yellow), MREs without seed match (grey) or genes without MREs (black). Numbers of transcripts in each set are given. Asterisks indicate the significance of difference to genes without MREs (ns>0.05, *<0.05, **<0.01, ***<0.001, asymptotic two-sample Kolmogorov-Smirnov test). **(e)** Changes in TE for transcripts with MREs vs. non-targets shown as $log_2$ fold-changes of KO over WT vs. density. Comparison of transcripts with different MREs as in (d). Numbers of transcripts in each set are given (ns>0.05, *<0.05, **<0.01, ***<0.001, Welch two sample t-test). **(f–i)** Comparison of transcripts with MREs with the canonical AA seed match sequence (yellow), the alternative AU seed match (red) for 6mer-m8 (f), 6mer (g), 7mer-A1 (h) and 7mer-m8 (i), MREs without seed match (grey) or genes without MREs (black) (ns>0.05, *<0.05, **<0.01, ***<0.001, asymptotic two-sample

Kolmogorov-Smirnov test). Only transcripts with a single MRE were used in all analyses shown.

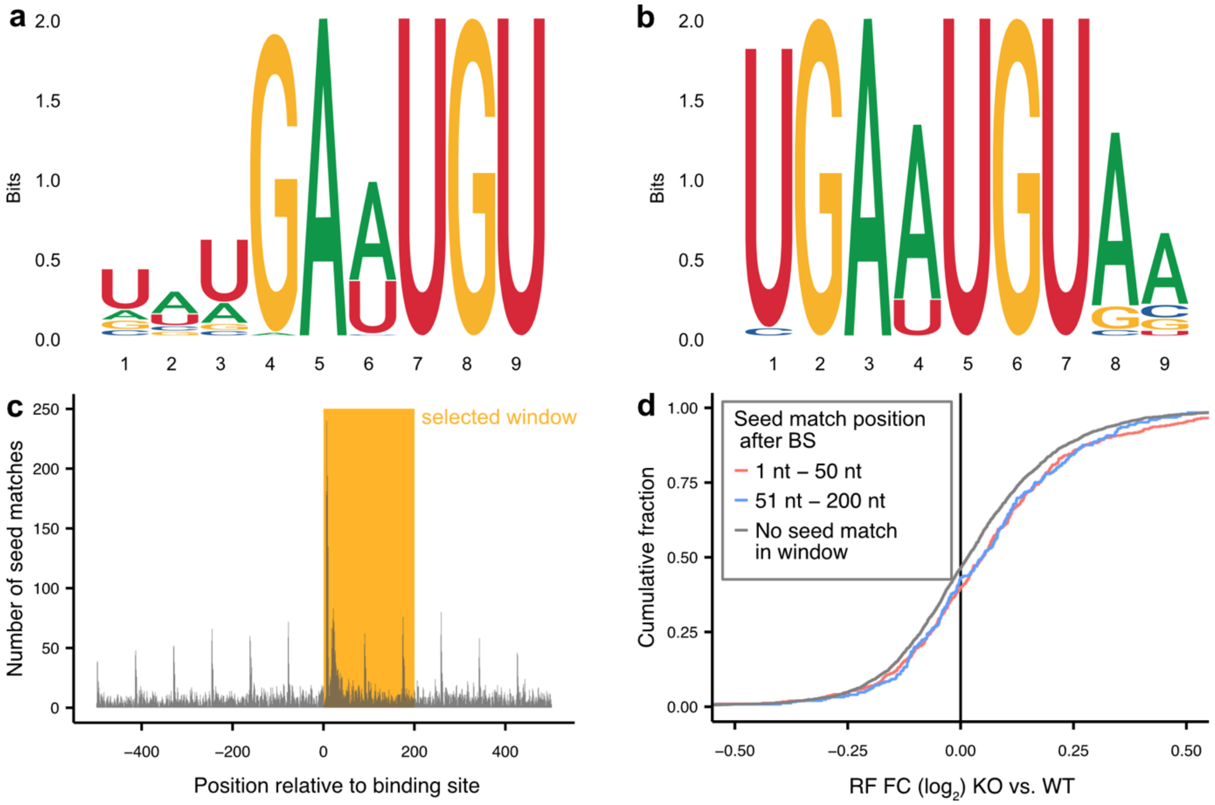


**Figure S8: Predominant sequence motifs and seed match identification are largely independent of the size of search windows around Ago2 binding sites. (a,b)** XSTREME top motif within a window of 101 nt (a) or 1001 nt (b) centered on the miR-181 binding sites. **(c)** miR-181 seed match positions relative to Ago2 binding sites. The orange box indicates the window selected for downstream analyses. **(d)** Fold changes in ribosome occupancy between miR-181 KO and WT conditions are shown as the cumulative density of the $\log_2$ values. Comparison of transcripts with MREs containing seed matches within 1 and 50 nt (red) and 51 and 200 nt (blue) downstream of the Ago2 binding site.

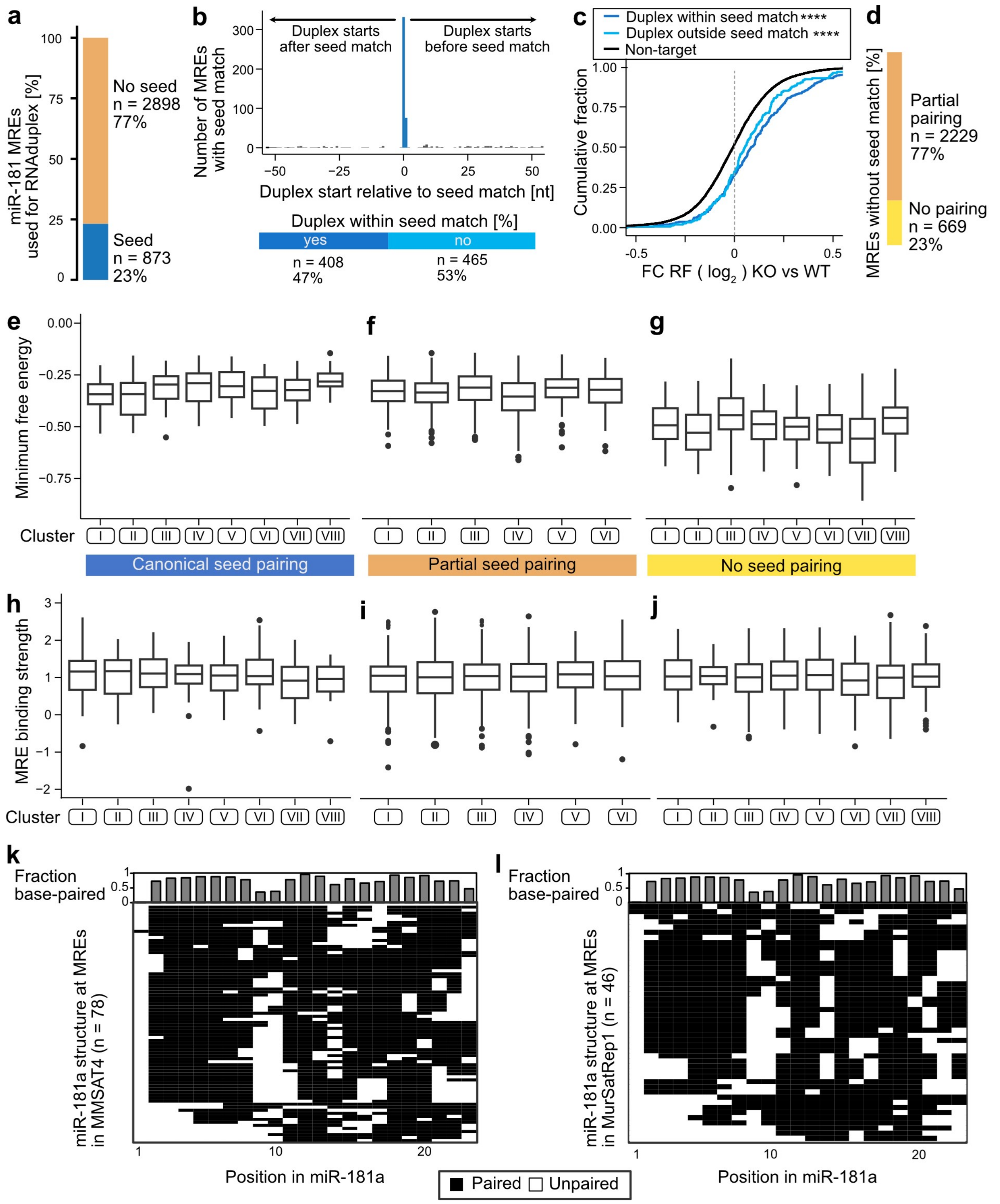


**Figure S9. Prediction of miR-181 pairing with target MREs. (a)** Ratio of miR-181 MREs used for RNAduplex containing a seed match (blue) or not (orange). **(b)** Top: Histogram of duplex start position relative to the start of seed match in MREs with seed match. Shown is a window spanning 50 nt before to 50 nt after the start of the seed match. Bottom: Ratio of duplexes starting at the start of the seed match or one nt before (dark blue) to duplexes starting at other positions (light blue). **(c)** Fold changes in ribosome occupancy between miR-181 KO and WT conditions are shown as the cumulative density of the $log_2$ values. Shown are duplexes starting at the start of the seed match or one nt before (dark blue), duplexes starting at other positions (light blue) and genes without MREs (black). Asterisks indicate the significance of difference to genes without MREs (****< 0.001, Kolmogorov-Smirnov test).

**(d)** Ratio of miR-181 MREs without a seed match with either partial (orange) or no (yellow) pairing within the first 8 nts of the miR-181a sequence. **(e–g)** Boxplots of the normalized minimum free energy per cluster for canonical duplex formation (e), partial seed pairing (f), or no seed pairing (g). Center line depicts the median, box encloses the 25% and 75% quantiles. Whisker is the maximum value of the data that is within 1.5 times the interquartile range over or under the 75th (upper) or 25% (lower) percentile respectively. **(h–j)** Boxplots of the MRE binding strength per cluster for canonical duplex formation (h), partial seed pairing (i), or no seed pairing (j). Center line depicts the median, box encloses the 25% and 75% quantiles. Whisker is the maximum value of the data that is within 1.5 times the interquartile range over or under the 75th (upper) or 25% (lower) percentile respectively. **(k, l)** Heatmap of miR-181a structures in duplexes with targets overlapping MMSAT4 (k) or MurSatRep1 (l) repeats. White – unpaired, black paired. Bars on the top annotate the paired ratio per nucleotide.
